# Single cell transcriptomes reveal characteristics of miRNA in gene expression noise reduction

**DOI:** 10.1101/465518

**Authors:** Tao Hu, Lei Wei, Shuailin Li, Tianrun Cheng, Xuegong Zhang, Xiaowo Wang

## Abstract

Isogenic cells growing in identical environments show cell-to-cell variations because of stochastic gene expression. The high level of variation or noise could disrupt robust gene expression and result in tremendous consequences on cell behaviors. In this work, we showed evidence that microRNAs (miRNAs) could reduce gene expression noise in mRNA level of mouse cells based on single-cell RNA-sequencing data analysis. We identified that miRNA expression level, number of targets, targets pool abundance and interaction strength of miRNA with its targets are the key features contributing to noise repression. MiRNAs tend to work together as cooperative sub-networks to repress target noise synergistically in a cell type specific manner. Using a physical model of post-transcriptional regulation, we demonstrated that the accelerated degradation with elevated transcriptional activation of miRNA target provides resistance to extrinsic fluctuations. Together, through the integration analysis of single-cell RNA and miRNA expression profiles. We demonstrated that miRNAs are important post-transcriptional regulators for reducing gene expression noise and conferring robustness to biological processes.

## Background

Variations of gene expression are often caused by genetic or environmental diversities [1]. However, even genetically identical cells growing in same environments may display diverse phenotypes. Gene expression noise has been suggested as a major factor of such cell-to-cell variations. On the one hand, gene expression noise is significant to cell development and population evolution [1, 2]. Such variability can increase overall fitness in evolution by expanding the range of phenotypes [3]. On the other hand, noise can result in non-reproducibility coordinate cellular functions during tissue morphogenesis and homeostasis [4, 5]. Maintaining robust output in fluctuant environment is necessary condition for cells to function physiologically. Although stochasticity is inevitable in the process of gene expression, it has been found that most genes in mammalian cells show inapparent randomness to response to the cellular state changing and microenvironment fluctuation [6], how cells and organisms maintain the fidelity of gene expression has still not been fully understood.

As an indispensable part of post-transcriptional regulation, microRNAs (miRNAs) have been considered as important regulators in fundamental cellular pathways and organismal development processes [7]. Each type of miRNA is predicted to regulate dozens or even hundreds of target gene species in mammalian cells, but only a small portion of these targets is moderately repressed (rarely exceeds 2-folds) [8]. That is to say, in most cases, miRNAs are not functioned as intensive repressors. Why are there so many evolutionary conserved miRNAs and potential targets if miRNAs are inefficient regulators? One proposed assumption is that miRNAs can modulate variations of gene expression and confer robustness of cell population phenotypes [7, 9-11]. Several recent works have studied the effect of miRNA regulation on gene expression noise in protein levels using synthetic gene circuits [9, 10, 12, 13]. Siciliano et al. [10] demonstrated that miRNAs provide phenotypic robustness to transcriptional regulatory networks by buffering fluctuations in protein levels. Schmiedel et al. [9] showed that miRNAs can reduces target gene intrinsic noise but with the cost of introducing extra noise of miRNA itself (miRNA pool noise, the fluctuation leaded by the changing of miRNA expression level [9]). Martirosyan et al. [13, 14] showed that the co-regulated targets of one miRNA species (competing RNAs) could enhance target stability across the full range of protein expression by buffering miRNA pool noise. However, whether these conclusions from synthetic gene experiments are applicable to endogenous genes is not clear. Furthermore, gene expression noise in protein level shows significant difference compared to mRNA level because of the translational bursting and the coupling between transcription and translation procedure [15-17].

The effects of miRNA regulation on mRNA expression noise using endogenous gene expression data is still lacking. On the one hand, there are dozens or even hundreds of types of miRNAs expressed in one single mammalian cell and each miRNA can have hundreds of target genes, which form a complex regulatory network [18, 19]. To characterize such direct or indirect interactions between targets in large interconnected networks, one can hardly study a single miRNA-target pair in isolation, but need to profile mRNA and miRNAs genome widely in a global view. On the other hand, high-throughput quantitative measurement of gene expression at single cell level were difficult in the past decade. smFISH and fluorescent reporter protein have been widely used to study gene expression noise [2, 20]. However, these approaches limit the number of genes that can be studied simultaneously, thus are not sufficient to provide a global picture to understand miRNA noise reduction. Development of single-cell RNA sequencing and its data analysis methods recently make it possible to evaluate RNA expression noise across cells in genome wide.

To better understand the influences of miRNAs on gene expression noise in mammalian cells, we combined single cell RNA-seq data with miRNA expression data to reveal pivotal features of miRNA mediating gene noise reduction that could not be oberserved by bulk measurements. We showed that miRNA reducing gene expression noise is a common property in various types of mouse cells. We found that miRNA target number, miRNA target pool abundance (the sum of target transcript counts), miRNA expression level, and miRNA-mRNA interaction strength are positively correlated with the strength of noise reduction, and miRNAs usually work together as co-regulation de-noising sub-networks to repress target noise synergistically in a cell type specific manner. A kinetic model was built to interpret the mechanism of miRNA reducing mRNA expression noise, which demonstrated that accelerated miRNA target degradation rate and elevated target transcription rate could contribute to the resistance of extrinsic fluctuations in environment, and the large competing target pool of miRNA could buffer miRNA pool noise. Our results suggested miRNAs are crucial for mRNA expression noise reduction and provided a new perspective to understand the physiological functions of miRNAs and their synergistic networks in vivo.

## Results

### miRNAs can suppress gene expression noise at genome wide

Accurate characterization of gene expression level is the precondition for studying expression noise. Single-cell RNA sequencing (scRNA-seq) provides a high-throughput measurement for acquiring the gene expression heterogeneity across cells [21]. However, current scRNA-seq measurements are suffered from non-negligible technical noise from stochastic mRNA loss, non-linear amplification and others variations in library preparation and sequencing [22, 23]. Therefore, unique molecular identifier (UMI) and spike-ins are recommended to control the level of technical variations [21, 24]. Here, three scRNA-seq datasets using both external RNA spike-ins and UMI from different cell types in mouse were used in this analysis (see Methods for details). We established a systematic data processing pipeline (Fig. S1) to eliminate technical variations and revealed the influences of miRNAs on gene expression noise. First, the true biological variations of UMI-based counts were separated from the high level of technical variations with the help of spike-ins [22]. Second, gene expression noise was quantified by the coefficient of variation (CV). The dependency of CV magnitude on gene expression level was removed by calculating local CV ranks in a sliding window of genes with similar expression levels. Then, we tested whether gene set that were targeted by miRNA had significantly lower expression noise than non-miRNA-target genes. The disparity between miRNA targets and non-targets was measured by the effect size of Mann-Whitney U test [25]. Finally, the effect size was corrected, named as AES score (adjusted effect size), to eliminate the influence of sample size so that we could compare the effect of noise reduction for different miRNAs (see Methods for details, Fig. S9-10). An AES score large than 0 means that miRNA target sets tend to have lower expression noise than non-targets, and vice versa.

We found that miRNA targets tend to have lower expression noise than non-targets in mRNA level (Fig. 1A). We identified target sets of 149, 108 and 30 miRNAs with significant positive AES score in mouse ES cells, intestinal cells and dendritic cells respectively, but none with significant negative AES in any of these three datasets (Fig. 1B). Such disparities were absent when using control samples generated by random sampling (Fig. 1C and 1D), which indicated that our noise analysis procedure could reveal true biological difference associated with miRNA regulation but not random technical variations in scRNA-seq. These observations suggested that miRNA reducing mRNA expression noise is a common feature across different cell types.

**Fig.1.**
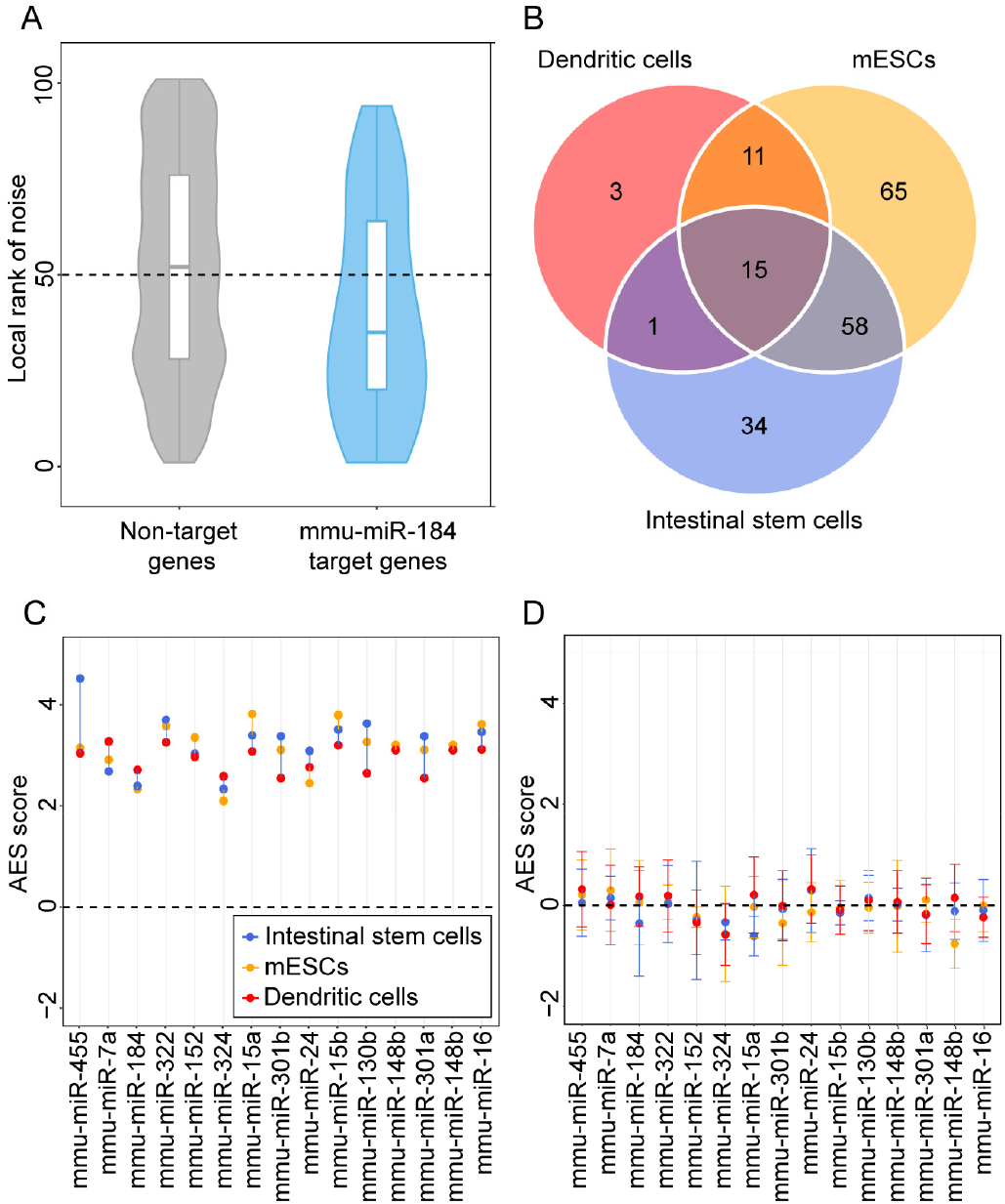
Gene sets noise analysis reveals that miRNA can suppress gene expression noise. (A) Box-violin-plots quantify the relative noise disparity of miR-184 target genes and non-miRNA-target genes. The middle line in the box is the median and the density represents the distribution of local rank of noise. The p-value is 1.9e-5 and AES score is 2.73 for statistic of Mann-Whitney U test. AES score of mmu-miR-184 large than 0 indicates that the expression noise of its target genes is lower than that of non-miRNA-targets. (B) Venn diagram shows the number of overlapped and specific de-nosing miRNA after multiple test correction (AES score > 0 AND false discovery rate (FDR) < 10%) among the three different cell types datasets. Colors indicate different cell types. (C) Noise disparity between the target genes and non-target genes for common de-noising miRNAs of three different cell types. Each data point is shown in three datasets and colors indicate different cell types. (D) Noise disparity are absent for random gene sets. As contrast test, the results of an identical analysis to that shown in (C), but instead using random samples as test and background. Points and error bars indicate mean and standard deviation over 100 samplings.

### miRNAs with large target pool could suppress noise better

Although miRNA target genes tended to be less noisy than non-target genes, there are still considerable differences in the noise levels among different miRNAs targets (Fig. 1C). This raised the question of what properties of miRNA may influence target mRNA expression noise. We tried to explore the major influence factors related to miRNAs noise reduction. We examined all 317 miRNAs expressed in mESCs [26] and found a significant positive correlation (P-value < 2.2×10^−16^, R-squared = 0.322) between miRNA’s target gene number and its effect on reducing gene expression noise (Fig. 2A). A positive correlation between the abundance of targets pool (the sum of the target transcript counts) and AES scores was also observed (P-value < 2.2×10^−16^, R-squared = 0.363) (Fig. 2B). An intuitional statistical measure of the relationship between high abundance of targets pool and noise reduction efficiency can be obtained by dividing the miRNAs into quadrants by AES score and targets pool abundance (Fig. 2B). Strikingly, when targets pool abundance is low, the fraction of miRNAs with low AES score is 7 times larger than when targets pool abundance is high (165/271=61%, 4/46=8.7%; odds ratio = 16.22, P-value = 8.9×10^−12^, Fisher’s exact test). Similar correlation can also be found in the relationship between high number of targets and high effects of noise reduction (odds ratio = 4.37, P-value = 3.1×10^−6^, Fisher’s exact test, Fig. 2A). These observations were also consistent in intestinal stem cells (Fig. S11) and dendritic cells (Fig. S12).

**Fig.2.**
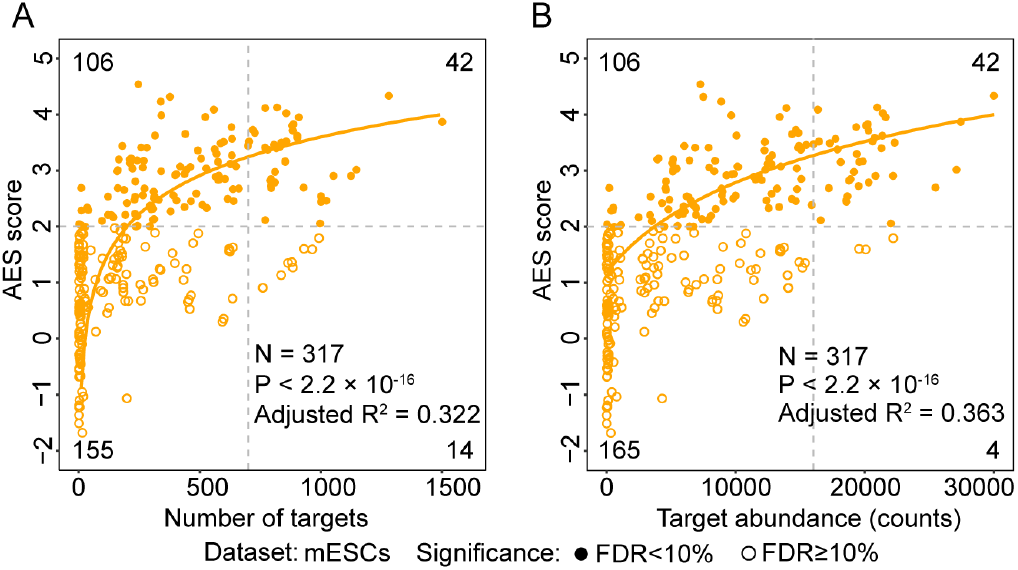
miRNAs with large target pool are associated with better noise suppression. (A) AES score of Mann-Whitney test versus number of targets across all detected miRNA in mESCs. Curves were fit to *a* + *b*log_10_(*x + c*) where a, b and c were determined by least squares error. Fitted parameters are shown in Table.S1. The cutoff for AES score is 2.0 and cutoff for number of targets is 700 in Fisher exact test. (B) Similar points and fitting curve to those shown in A for targets pool abundance. The cutoff for AES score is 2.0 and cutoff for targets pool abundance is 16000.

Taken together, miRNAs with large target pool tend to suppress target gene expression noise better.

### miRNA-target interaction strength influences noise reduction effect

miRNAs can bind to the miRNA response elements (MREs) in their target mRNAs [27]. The binding strength between miRNA and its target mRNA can be quantified using RNA hybridization free energy in thermodynamic which is correlated with the miRNA’s ability to repress translation of the mRNA [28]. In order to study the relationship between noise reduction and miRNA-mRNA interaction strength, we collected free energy information from miRmap [29] for all possible miRNA-mRNA pairs in mouse. To compare their relative binding energy strength among different miRNAs-target pairs, all the miRNA-target pairs were ranked in an ascending order according to their binding energy [29], and were then divided into three sets based on quantiles. The top 40 percents pairs (low free energy) indicates a strong interaction between miRNA and mRNA, while the bottom 40 percents pairs (high free energy) indicates a weak interaction (Fig. 3A). The effects of noise reduction for this two sets were examined respectively, and the results showed that strong interactions will enhance the effect of noise reduction in all test datasets (Fig. 3B for mESCs, Fig. S13A for intestinal stem cells, Fig. S13B for dendritic cells), which is concordant with previous analysis that miRNA reduces intrinsic noise of gene expression in protein level by the miRNA mediated fold repression [9].

**Fig.3.**
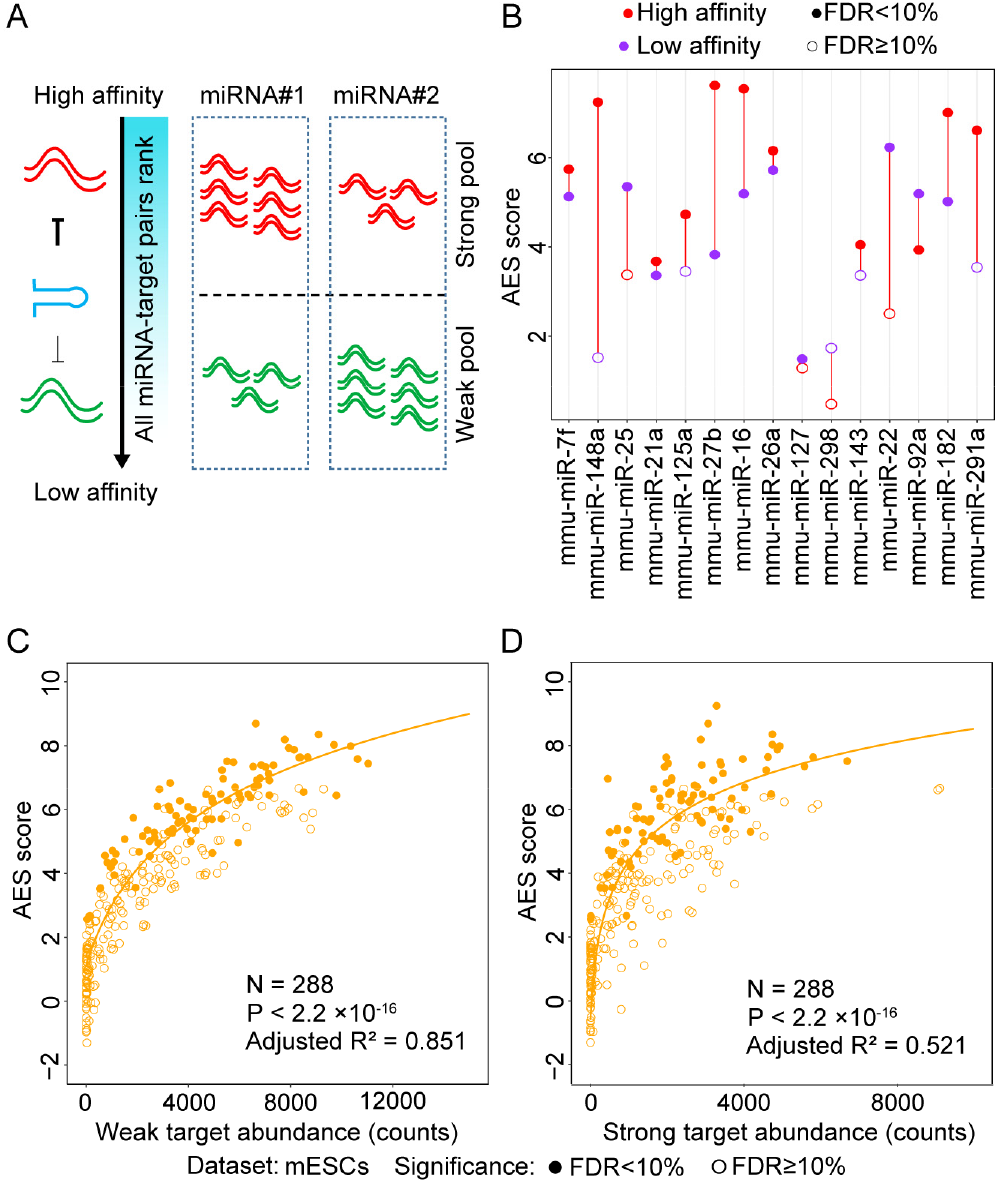
miRNA-target interaction strength influences noise reduction effect. (A) The binding energy of all miRNA-mRNA pairs are ranked in ascending order. The endogenous miRNA target pool is indicated in wavy lines illustrating the targets affinity (red for high affinity, green for low affinity) and their relative abundances. (B) Noise disparity between the target genes and non-target genes of top15 abundant miRNAs for strong (red) and weak interaction strength (purple) in mESCs respectively. (C-D) AES score versus weak targets pool abundance and strong targets pool abundance across all miRNAs expressed in mESCs respectively. Curves were fit to *a + b*log_10_(*x + c*) where a, b and c were determined by least squares error. Fitted parameters are shown in Table. S1.

We further explored the contribution of different interaction competing targets pool abundance to the noise reduction effect for the strong interaction targets (Fig. 3C and 3D). Though both the strong and weak competing target pool size were positively correlated to the AES score, the predictive power of the weak competing target pool size (regression R-squared = 0.851) were much higher than the strong competing target pool size (regression R-squared = 0.521). These observations were consistent with our previous theoretical research on miRNA regulation noise [14], that abundant weak competing target RNA pool has the capacity to buffer gene expression noise.

Taken together, a gene strongly bounded by miRNAs, which have large weak interaction competing target pool, is tended to have lower mRNA expression noise.

### Gene expression noise are repressed by miRNA co-regulation sub-network

Intuitively, we expected that the miRNA expression level should have an influence on noise reduction. However, we did not find a strong correlation between the miRNA expression level and the AES score (Fig. 4B). One possible reason is that each miRNA species can regulate multiple target genes and each gene can be controlled by multiple miRNAs, which constitute a complex regulation network. The common targets of different miRNAs could induce indirect miRNA-miRNA crosstalk [30], which may mask actual interaction of miRNA with targets and lead to false result in AES score calculating. What’s more, miRNAs seldom regulate cellular processes independently but often construct functional miRNA–miRNA cooperation networks via co-regulating functional modules [31, 32]. To study miRNA cooperative regulation and analyze the function of miRNA in noise reduction in sub-network level, we predicted miRNA-miRNA cooperation network through their common target genes. Each miRNA species was regarded as a node of the network, and the weights of the edges between the nodes were calculated by the similarity of their target gene sets. Walktrap clustering algorithm [33], which is suitable for the subnetwork division problem of complex networks, was performed on the 317 expressed miRNAs in mESCs and identified 59 sub-networks (Fig. 4A). The AES score of noise reduction was calculated for the sub-networks, by taking the targets of the miRNAs in the sub-network as a union. As shown in Figure 4, AES scores of the miRNA clusters were highly correlated with miRNA expression level (Fig. 4C), number of targets (Fig. 4D) and target abundance (Fig. 4E). Similar results were also obtained in intestinal stem cells (Fig. S14) and dendritic cells (Fig. S15). Importantly, such strong positive correlations are much higher than those measured in the level of miRNA individuals or miRNA families defined by miRNA 5’ end 2-8nt ‘seed’ similarity (Fig. 4F). These observations by clustering analysis successfully revealed biological significant noise reduction by miRNA sub-networks, as well as the strong correlation between noise reduction with number of targets, abundance of targets pool and miRNA expression measured in sub-network level.

**Fig.4.**
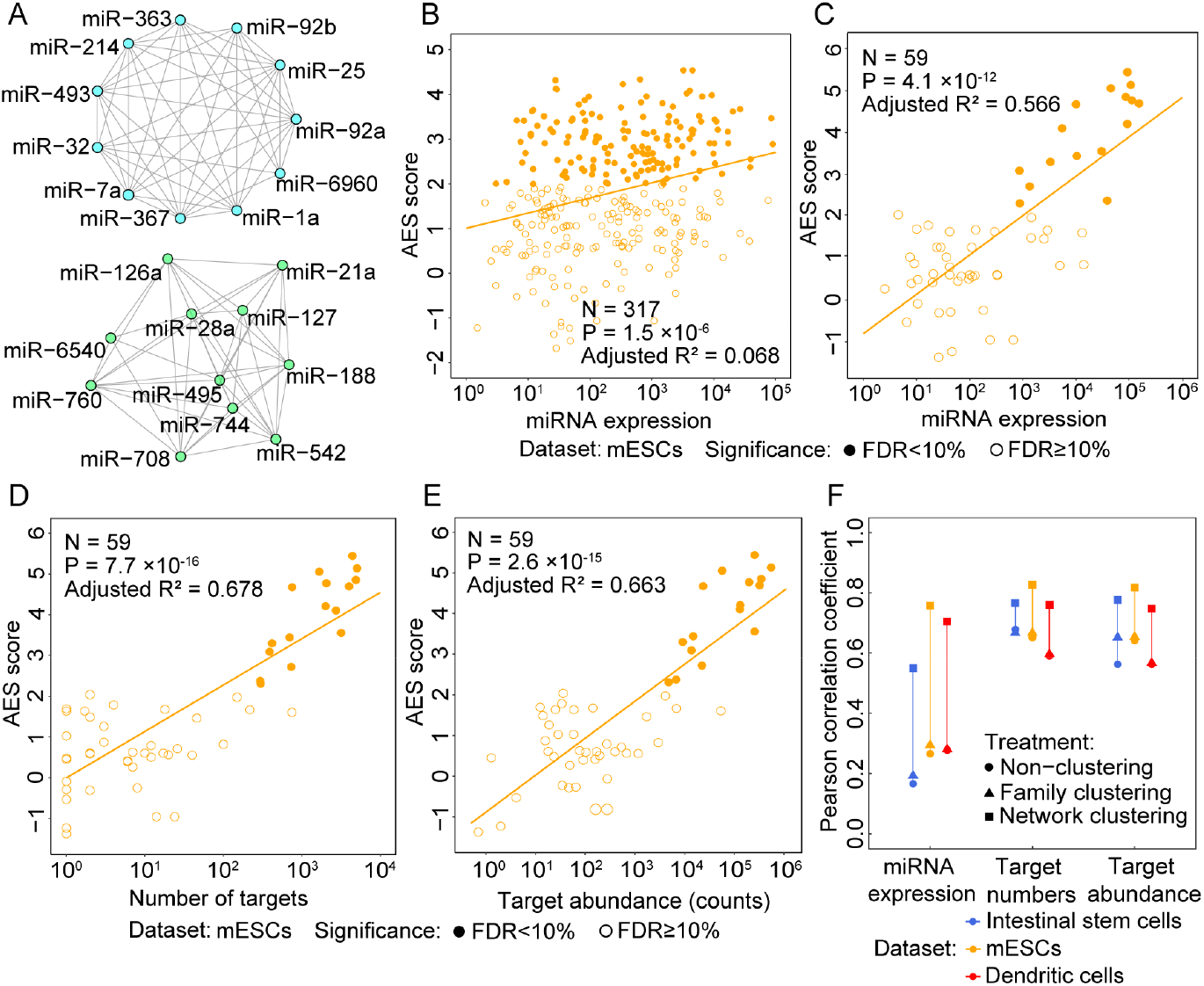
Gene expression noise are repressed by miRNA co-regulation sub-network. (A) Two examples of significantly de-noising miRNA sub-networks for mESCs. Nodes represent miRNAs and edges indicate the connection of miRNAs. Others significantly de-noising miRNA sub-networks are shown in Fig. S16 and Table. S4. (B-C) AES score versus miRNA expression for miRNA individuals and sub-network respectively. Curves were fit to *a* + *b*log_10_*x* where a and b were determined by least squares error. Fitted parameters are shown in Table. S2. (D) AES score after sub-network clustering versus number of targets across all cluster miRNA in mESCs. (E) Similar points and fitting curve to those shown in C for targets pool abundance. F) Pearson correlation coefficients between miRNA feature (log transferred) with AES score for non-clustering (circle), clustering by miRNA family (triangle), and clustering by network (square) respectively. Colors indicate different cell types.

### miRNA co-regulation sub-network showed cell-type specificity

We next compared miRNA de-noising sub-networks in different cell types. In total, 37 significantly de-noising miRNA subnetworks were identified in three cell types (16 in embryonic stem cells, 11 in intestine stem cells, and 10 in dendritic cells). For any two given miRNA sub-networks, we first obtained their target genes sets A and B, and then calculated sub-network similarity using Jaccard similarity coefficient [34] of their target genes (defined as the intersection size divided by the union size of two target genes sets), and then constructed a similarity matrix for all miRNA sub-networks from three cell types. These sub-networks were divided into constitutive sub-networks and cell-type-specific sub-networks by hierarchical clustering of the similarity matrix (Fig. S16). The constitutive sub-networks regulate common target sets across three cell types and the cell-type-specific sub-networks tend to target specific genes in a given cell type. To further investigate the roles of these two types of sub-networks, gene ontology analysis was performed (see Methods for details) (Fig. 5). The target genes of constitutive de-noising miRNA sub-networks were enriched in terms related to essential cellular function such as cytoplasmic mRNA processing body assembly, protein catabolic process and ribonucleoprotein complex assembly. In contrast, cell-type-specific miRNAs sub-networks might regulate more specific biological processes such as body morphogenesis in mouse embryonic stem cells, glycoprotein metabolic process and epidermal growth factor receptor signaling pathway in dendritic cells, cell cycle phase transition and mitotic cell cycle phase transition in intestinal stem cells, which implied that cell-type-specific miRNA sub-networks may contribute to cell state maintenance by repressing cell-type-specific gene expression noise. In summary, these observations suggested that miRNA subnetworks, which correspond to cell types and cell functions, play important roles in gene expression noise reduction.

**Fig.5.**
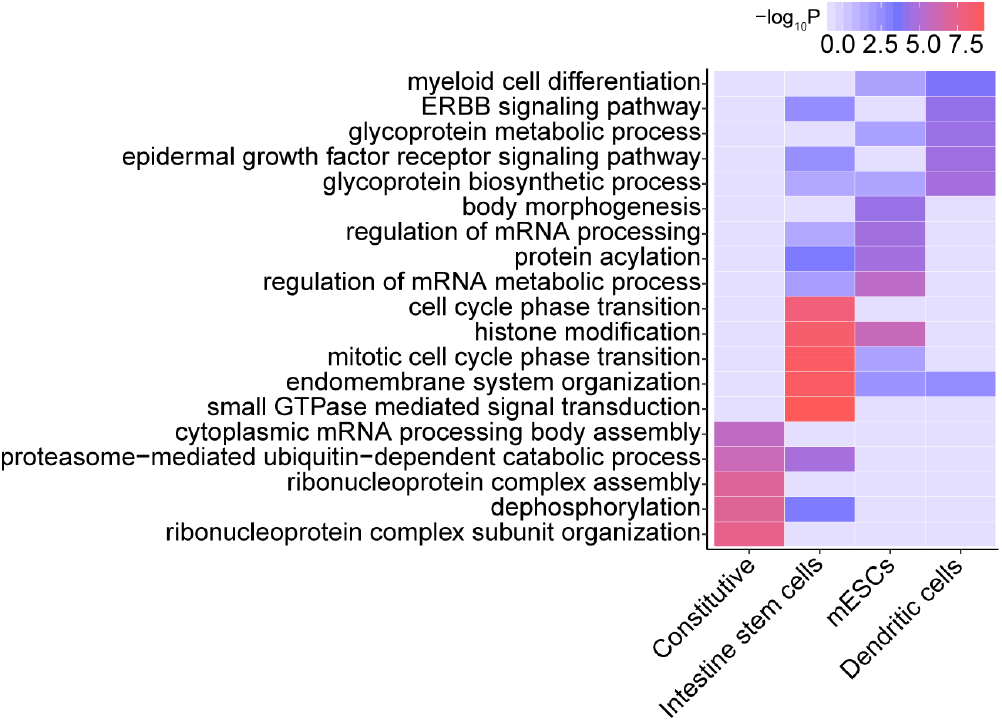
miRNA de-noising subnetworks are corresponding to cell types and cell functions. Top 5 enriched gene-ontology (GO) terms with their respective p values (clusterProfiler) for target genes in constitutive de-noising miRNAs sub-network and cell-type specific de-noising miRNA sub-network in mESCs, intestine stem cells and dendritic cells respectively.

### Accelerated miRNA target degradation provides resistance to extrinsic noise

The half-life period of mRNA is known to influence gene expression noise [35, 36]. Due to the integral effect on transcriptional bursts, longer-lived mRNAs generally exhibit lower expression intrinsic noise. With the analysis combining the mRNA degradation rate data [37] with scRNA-seq data, the trend can be clearly observed that mRNA half-life is negatively correlated with mRNA expression noise (Fig. 6A for mESCs, Fig. S17A-B for intestinal stem cells and dendritic cells). However, miRNAs usually repress gene expression by accelerating target mRNA degradation [9]. The half-life and noise of genes targeted by miRNAs exhibited a counter-intuitive positive correlation (Fig. 6A, Fig. S17A-B). Interestingly, the target genes of de-noising miRNAs have even significantly shorter half-life than those targets of non-de-noising miRNAs (Fig. 6B for mESCs, Fig. S18A for intestinal stem cells and Fig. S18B for dendritic cells). Recently, Schmiedel et al. found that miRNA target genes usually show compensatory elevated transcriptional activation to compensate miRNA repression [38]. Therefore, one conjecture for this seemingly contradictory phenomenon is that the target genes of miRNAs resist extrinsic disturbances and maintain robust gene expression through a higher transcription and degradation rates.

**Fig.6.**
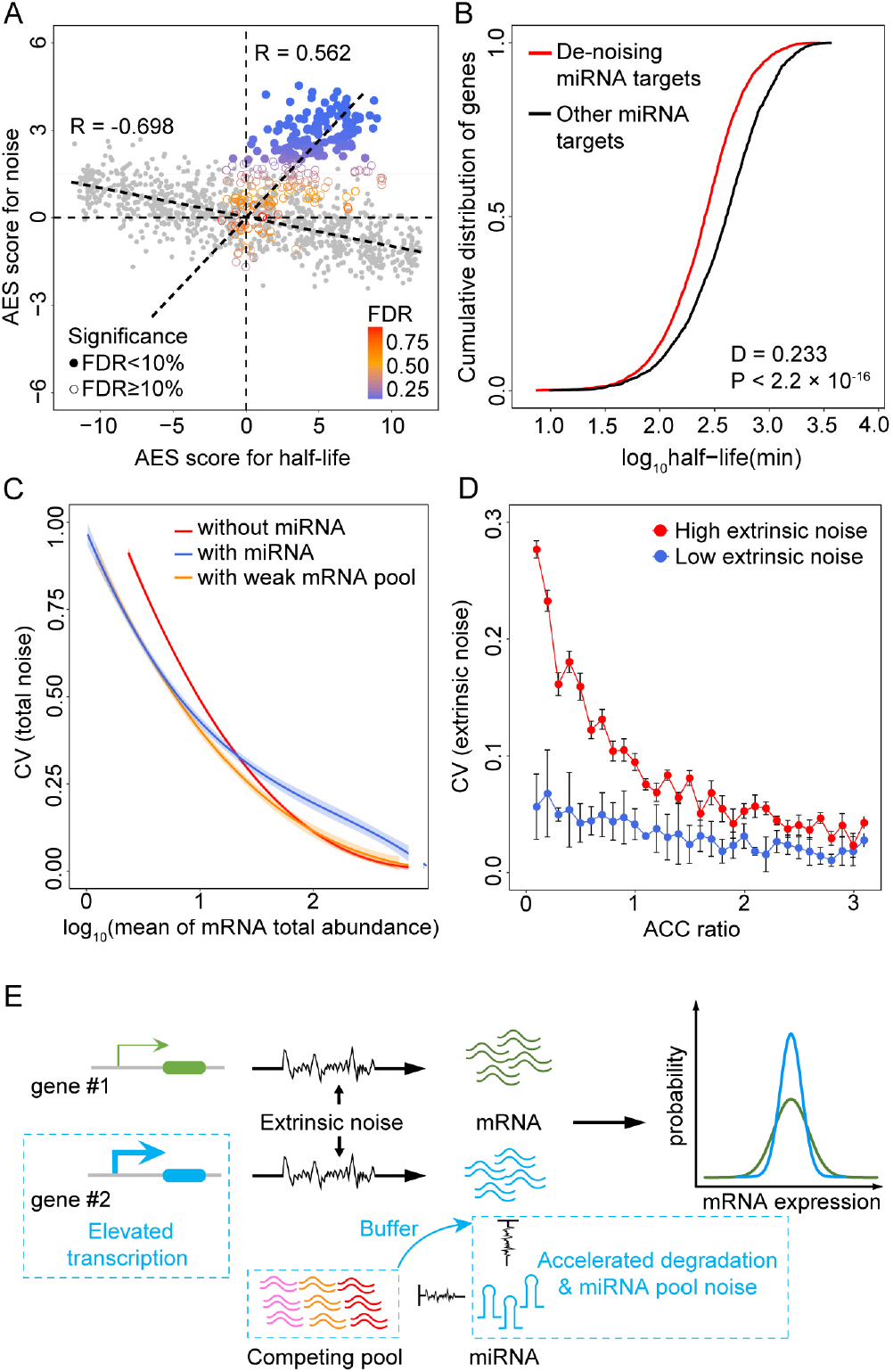
miRNAs accelerate target degradation to resist environmental disturbances. (A) Scatterplot of gene expression noise AES scores versus mRNA half-life AES scores for random gene sets (grey) and real miRNA target gene set (color). Random gene sets are constructed with varying sample size (100–2000) by weighted sampling to mRNA according to half-life. AES score is defined as the adjusted statistic for Mann-Whitney U test, where a value of 0 corresponds to no difference between gene sets and background while a value large than 0 correspond to a lower noise and lower half-lives gene sets than background. The Spearman correlation coefficient between half-life and noise AES score for random sample is −0.698 but 0.562 for miRNA target genes in mESCs. Similarity results obtained using intestinal stem cells and dendritic cells datasets are shown in Fig.S17A-B. (B) Targets of de-noising miRNAs have significant shorter half-life than those targets of non-de-noising miRNAs in mESCs. (C) Total noise (intrinsic and extrinsic noise) of mRNA as a function of mean mRNA total expression (Sum abundance of free mRNA and mRNA-miRNA complex). The shadows indicate 100 repeated simulation trials. (D) The extrinsic noise of miRNA-regulated genes decreases as the accelerated degradation increases. The error bars indicate the standard deviation of 100 repeated simulation trials. (E) Summary scheme showing accelerated degradation with compensatory elevated transcriptional activation of miRNA target provides resistance to extrinsic fluctuations. And the large competing targets pool with weak binding affinity could buffer miRNA pool noise. Therefore, miRNA target gene (blue) have lower noise than non-target gene (green) at equal mRNA expression levels globally.

We established a coarse-gained model to investigate this hypothesis. The model described the main steps of gene expression, miRNA post-transcriptional regulation, and competing target regulation by miRNA, as illustrated in Fig. S2. It was inspired by our and others’ previously studies on stochastic gene expression and post-transcriptional regulation by miRNA [9, 39]. The noise in gene expression comes from two major sources: one is intrinsic noise generated by random biochemical reaction such as mRNA and miRNA production and decay, association and dissociation of free mRNAs with miRNAs; and the other one called extrinsic noise is modeled as the fluctuation in the reaction kinetic rates generated by variable external environment factors in cellular components such as the number of RNAPs or ribosomes [2, 40, 41]. We defined the strengths of extrinsic noise as Fano factors of reaction kinetic rates as previous studies [40] and extend the chemical Langevin equation to include both intrinsic and extrinsic noise (see Supplementary Materials). Without loss of generality, we assumed that the extrinsic noise has the same effect on all parameters, that is, all reaction kinetic rates share an equivalent Fano factor.

Model simulation suggested that, miRNA could reduce mRNA expression noise in the low expression zone but introduce extra miRNA pool noise in high level of mRNA expression compared to an unregulated gene at equal mRNA expression levels (yellow line in Fig. 6C) similar to previous experimental results in protein level [9]. Keeping the level of miRNA repression to target gene unchanged, introducing large competing target pool with weak binding affinity could significantly buffer the noise introduced by miRNA pool (blue line in Fig. 6C). To explore the influence of accelerated degradation on noise, accelerated degradation (ACC ratio) is defined as the ratio of miRNA-induced mRNA degradation rate to mRNA degradation rate (see Supplementary Materials). Under the condition of equal level of mRNA expression, the model predicted that miRNA targets have reduced extrinsic noise when accelerated degradation degree increasing (Fig. 6D).

In summary, these observations indicated that accelerated target mRNA degradation with elevated transcription rate may contribute to the resistant of extrinsic fluctuations, and the large competing targets pool with weak binding affinity could buffer miRNA pool noise (Fig. 6E).

## Discussion

In this work, we revealed the miRNA regulation effect on target mRNA expression noise by using single cell RNA sequencing data in mouse. We showed that reducing target expression noise is a common feature for miRNAs in various mouse cells. We identified miRNA characteristics contribute to noise repression, including number of targets, targets pool abundance, miRNA expression level and interaction strength of miRNA with its targets. Furthermore, we showed that miRNA co-regulation subnetworks are significantly correlated with noise reduction, which is helpful to understand the function of miRNA co-regulation network in vivo.

We found that de-noising miRNAs targets have significantly shorter half-life than those targets of non-de-noising miRNAs. Based on these observations, we proposed that miRNAs, as noise suppressor, resist extrinsic fluctuation through increasing targets degradation rates with compensatory elevated transcription rates (Fig. 6E). In addition, we showed that the buffering role of miRNAs become evident with the increase of target degradation rates through kinetic model. Interestingly, Schmiedel et al. found miRNA targets half-lives decreased with the increase of number of microRNA sites but the transcription rates of targets increased with the increase of number of microRNA sites [38], which supported our assumption. Furthermore, there are several pieces of evidence which suggested miRNA could buffer gene expression noise against extrinsic environmental perturbation. Xin et al. found that miR-7 is essential to buffer developmental programs against variations and impart robustness to diverse regulatory networks, which was strongly functional conservation from annelids to humans [42]. Mehta et al. showed that miRNAs in hematopoietic system could modulate the balance between self-renewal and differentiation, and ensure appropriate output of immune cells, which conferred robustness to immune cell development, especially under conditions of environmental perturbations [43]. Kasper et al. found that miRNA mutant embryos of zebrafishes showed higher sensitivity to environmental perturbations and the loss of miRNAs increased the variance of developing vascular traits [44]. These evidence highlight the potential importance of miRNAs in stabilizing gene expression variability and preventing environmental susceptibility.

Previous studies were mainly focused on the role of miRNAs but neglected the miRNA-mediated cross-talk among competing targets. Recent works have found that the cross-talk among competing targets could enhance the stability of gene expression [13, 14, 45]. Here, we observed a positive correlation between the abundance of targets pool and noise reduction (Fig. 2 and Fig. 4). Interestingly, we found that weak targets pool has much larger capacity to buffer miRNA pool noise than the strong targets pool. MiRNAs are ubiquitous but mysterious regulators in mammalian cell, the majority of mRNAs are predicted as targets for one or more miRNA species, but only a small portion of those targets could be moderately repressed (rarely exceeds 2 fold change) [46]. Most miRNA-target interactions are too weak to have strong phenotypic consequences when knocking down or knocking out corresponding miRNA. The function of those pervasive and evolutionary conserved miRNA-mRNA weak interaction pairs is still unclear. Our results provided a possible direction to explain the weak repression effect exerted by the majority of the known miRNAs. In general, introducing higher level of miRNAs and compensable weak co-regulated targets could enhance the suppression of gene expression noise in a wider range.

This study focuses on the integrative analysis of scRNA-seq expression and bulk miRNA expression in various cell types. We selected the bulk miRNA expression data corresponding to the cell type of scRNA-seq from the public GEO database, and performed quality control to exclude the poor-quality miRNAs across different datasets. However, extensive genetic and expression variations of miRNA exist between different cells [47, 48], and the heterogeneity between cell individuals could make it difficult to find an accurate regulatory relationship between miRNA and their targets. Understanding endogenous miRNA functions precisely requires approaches to profile miRNAs and their targets simultaneous in single cell resolution. Though some approaches that enables high-throughput sequencing of single cell miRNAs were proposed very recently [49, 50], the paired high quality data of miRNA and their target in the same single cell are still lacking, which prevents for the discovery of associations between transcriptional and miRNA profile variation. These questions may be further addressed by multi-omics profiling of single cells in the future.

In summary, this work presented genome-wide evidence that miRNAs function as gene expression noise repressors, and the miRNAs function as networks to stabilize mRNA level and maintain cellular functions.

## Conclusions

Combining single cell RNA-seq data with miRNA expression profiles, we found consistent evidences of a role for miRNAs as noise repressor in multiple cell types of mouse. We systematically analyzed the key properties of miRNAs associated with noise reduction from individual miRNAs to miRNA co-regulated networks. Specifically, we proposed a kinetic model to reveal that increasing degradation rate to resist extrinsic fluctuations is the strategy for miRNAs to decrease mRNA expression noise. Gene expression processes are inherently stochastic under complex and volatile environment condition. However, the cellular mechanism control gene expression noise has not been well understood. Our results provide new insights to explain the role that miRNAs play when cells respond to environmental disturbances.

## Methods

### Expression quantification and normalization for scRNA-seq data

To systematically study the influences of miRNA regulation on gene expression noise in mammalian cells, we collected three single cell RNA-seq datasets employing external RNA spike-ins and unique molecular identifier (UMI) from different cell types in mouse, which consist of 41 cells from mouse embryonic stem cells [51], 90 cells from dendritic cells [52] and 100 cells from small intestinal stem cells [53] after removing low sequencing quality cells. We also collected bulk miRNA-seq data for the corresponding cell types [26, 54, 55]. UMI are tens of thousands of short DNA sequences incorporated in mRNAs before library amplification to account for the stochastic RNA loss and non-linear amplification bias for diversity expressed genes [21]. Spike‑ins are extrinsic molecules expected to be the same across all single-cell libraries and can be used to estimate technical variation in sequencing [22]. We collected single-cell RNA-seq datasets that were sequenced using UMI and spike-ins to reduce technical noise and estimate the actual biological signals used for downstream analysis. These features are very important to remove strong technical noise of scRNA-seq data to unveil subtle gene expression variation regulated by miRNA.

In order to compare expression noise between genes of different expression level and transcript length, the raw single-cell RNA-seq data were processed referring to the previous work [22]. First, to restore the true biological signal from the high level of technical noise coupling in the sequencing process of scRNA sequencing, we estimated gene expression mean and biological variance by accounting for technical noise with the help of spike-ins (Fig. S3-5) [22]. The scRNA-seq data analysis procedure based on UMI and spike-ins provides a reliable estimates of gene expression variation. This step controls various technical variation such as mRNA stochastically loss, amplification bias, sequencing efficiency variation, sampling variation and experimental batch [22] prior to downstream quantitative analyses. Second, we checked whether cell cycle relevant gene contribute substantially to the heterogeneity of gene expression. If so, the cell cycle co-factor would need to be removed out to ensure that it does not introduce spurious correlations or inflate the variances [53, 56]. As shown in Fig. S6-8, by comparing the variance distribution of cell cycle genes and other genes, cell cycle related genes do not show increased variability and are thus unlikely to lead to false results in the noise analysis.

### Gene expression noise quantification

We quantified gene expression noise by the coefficient of variation (CV, standard deviation divided by mean abundance of mRNA) (Fig. S3-5) as previous analyses [57, 58]. As CV is strongly anti-correlated with the expression level of genes (Fig. S3B-5B), the local ranks of CV in a sliding window of 100 similar expressed genes were calculated to remove the dependency of CV on gene expression level (Fig. S2). The local rank of CV is a reliable and robust measurement to compare relative noise level for gene sets with different expression [25].

### Noise disparity analysis

Referring to the previous work [25], we calculated the effect size to compare differences in noise level between miRNA targets and non-targets. Firstly, we retrieved miRNA expression level from miRNA-seq datasets, and the miRNA and mRNA interactions from TargetScan predictions [59, 60]. After that, we divided all genes into miRNA target gene sets and non-miRNA-target genes and compare the local rank of CV between the sets of target genes and non-miRNA-target genes. The significance of each test is determined using a Mann-Whitney U test, which is a non-parametric test that can be used in place of an unpaired t-test [61]. For each test, the null hypothesis is that the CV local rank of two genes sets come from the same population. In this context, we use the area under the ROC curve (AUC) statistic to measure effect size of the test [62] (Fig. S2). The AUC is equivalent to the Mann-Whitney U-statistic in mathematics [63]. Effect size great than 0.5 corresponds to higher expression noise of miRNA target genes than non-targets, and effect size less than 0.5 indicates lower expression noise of miRNA target genes than non-targets.

The effect size magnitude of Mann-Whitney test is correlated with the sample size (Fig. S9A). As different miRNAs have different number of targets, we need to correct the influence of sample size on the effect size to make them comparable between different miRNAs. Mathematically, effect size can be approximated as a normal distribution for large sample size, whose expectation *μ* is 0.5 and variance *σ*^2^ is 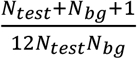, where *N_test_* and *N_bg_* represent the sample size of test and background respectively [63]. To remove the effects of sample size, the adjusted effect size (AES) was calculated: Adjust effect size = 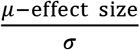. Fig. S9B shows AES score for different test sample size. The AES score less than 0 corresponds to a larger local rank CV of miRNA target genes than non-targets, and AES score large than 0 indicates that local rank CV of miRNA target genes are smaller than non-targets. Moreover, larger AES score means better noise reduction for miRNAs.

### Gene ontology analysis

To investigate the roles of constitutive miRNA sub-networks and cell-type specific miRNA sub-networks, we performed gene ontology analysis by genome wide annotation database org.Mm.eg.db and gene functional annotation clustering tool clusterProfiler with default setting [64, 65]. We chose the top 5 GO terms from constitutive miRNA sub-networks and each cell-type specific miRNA sub-networks and provided 19 terms in total. (Fig. 5, the Benjamini-corrected P value of these terms are less than 0.05).

### Model to describe the miRNA de-noising gene expression

A detailed description of theoretical analysis used in this study is available in SI Materials and Methods. Briefly, we built a physical model to investigate the mechanism of miRNA reducing gene expression noise in mRNA level.

### Additional software and code availability

Our analysis is based on R version 3.4.2. The codes used to perform the data process and analyses are available upon request.

## Abbreviations

scRNA-seq: single cell RNA sequencing
RNA-seq: RNA sequencing
miRNA: microRNA
mRNA: messenger RNA
UMI: unique molecular identifier

## Additional files

**Supplementary Materials. PDF** including all supplementary methods, Figure S1–18 and Table S1-4.

## Authors’ contributions

T.H, X.W and X.Z designed research, T.H, L.W, S.L and T.C performed analysis. T.H, X.W., L.W and X.Z wrote the manuscript. X.W. conceived and supervised this study. All authors read and approved the final manuscript.

## Acknowledgements

We thank Guiying Wu and Xianglin Zhang for technical assistance. We thank Prof. Jingzhi Lei for critical advice in the kinetic model.

## Supplementary Methods and Figures

### 1. The workflow of gene expression noise analysis

As shown in Fig. S1. The workflow for gene expression noise analysis consists of four main steps.

The first step is to quantify the expression of single cell transcripts and miRNA transcripts.

After that, we normalized the raw transcripts counts to remove the technical noise coupled in sequencing process with the help of Spike-ins [1]. And we quantified noise as coefficient of variation(CV) and calculated the local rank of CV in a sliding window to remove the dependency of CV on gene expression level as [2]. In the subgraph of second steps, left: coefficient of variation versus mean of transcript counts for all detected genes in raw data from scRNA-seq experiment in mESCs, black dots and red dots indicate true genes and spike-ins respectively. Middle: Predicted CV of transcript counts versus predicted mean transcript counts for all detected genes after removing technical noise with the help of spike-ins. Right: The rank normalization step removes the dependency of CV on gene expression level; that is to say, mapping the predicted CV to local rank of noise for every single gene.

In the third step, we predicted the interactions of miRNA and mRNA with TargetScan [3, 4]. Then we tested whether sets of genes that regulated by miRNA have unusually lower expression noise than non-target genes with Mann-Whitney U test and measured the effect size of test using the area under the ROC curve (AUC) statistic. In the subgraph of third steps, left: predicting the interactions of miRNA and mRNA with TargetScan. Middle: hypothesis test with Mann-Whitney U test. Right: using the area under the ROC curve (AUC) statistic which is equivalent to the Mann-Whitney U statistic to measure the noise bias of miRNA target genes.

In the fourth step, we corrected the effect size, named as AES (adjusted effect size), to eliminate the influence of sample size. The AES large than 0 means that the targets of miRNA have usually low expression noise than non-targets, and vice versa. Left: we validated the correlation of effect size and test sample size by random sampling background samples and variable test samples size (x-axis) from all genes in three cell types datasets. The three cell types datasets are highlighted with different color. And all tests are not significant in random experiment (FDR<10%). Meanwhile, this simulation result show that effect size is a strongly correlated with the test sample size. Right: An additional effect size correction step to remove the effects of sample size by calculating the adjusted effect size as z-score according the distribution of effect size. This plot shows the AES (adjusted effect size) which is independent to sample size after correction.

### 2. The coarse-gained kinetic model for miRNA de-noising gene expression

Here we modeled the noise of a microRNA regulated gene in the mRNA expression. This physical model was inspired by previously studies on stochastic gene expression and post-transcriptional regulation of miRNA [5, 6]. In brief, this physical model describes a sequence of biochemical reactions associated gene expression and post-transcription regulation, including the transcription and degradation of mRNA, the reversible binding of miRNA with mRNA which accelerates the degradation of mRNA and inhibits translation of the mRNA into protein. Parameters involved in the model where miRNA (*R*) regulate a target mRNA species (*m*) are described as follows. In general, *m* and *R* is produced with a rate of *k_m_, k_R_*, respectively. Free *m* and *R* degrades at a rate of *g_m_*, and *g_R_*. *m* binds to *R* to form complex *C* at a rate of *k_on_* or *k_off_*, and the complex *C* dissociates into *m* and *R* at a rate of *k_off_*. Complex *C* degrades at a rate of *gc* and *α* represents the miRNA loss rate (Fig. S2).

The stochastic gene expression model is described in the following ODEs:

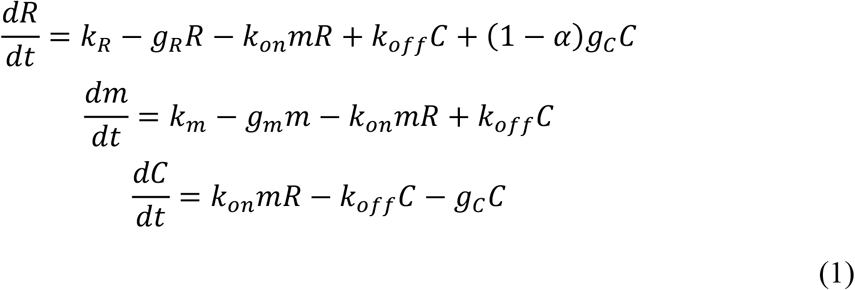

The biochemical reactions above are inherently stochastic because of the random collisions of the molecules. Considering a system of N molecules and M reaction channels, we can describe this random process with chemical master Eq. (3)

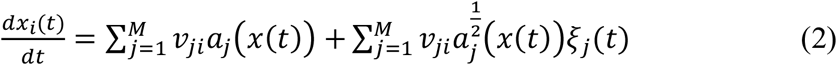

where *x_i_*(*t*) is the number of the *i*th molecules, *a_j_*(*x*(*t*)) is the propensity function of the *j*th reaction channel when *x* is equal to *x*(*t*), *ν_ji_* represents the amount of change in the *i*th molecule when the *j*th reaction channel occurs, the *ξ_j_*(*t*) is a Gaussian white noise with zero mean, time-independent, and statistically independent. Chemical Langevin equation only describes the intrinsic noise in gene expression. The other major noise resource called extrinsic noise is fluctuation in the reaction kinetic rates caused by variable external factors in environmental. Assume that the propensity functions depends on the reaction rates {*r*_1_, *r*_2_, *r*_3_.. *r*_*K*_}, and there is noise perturbation on the reaction rates, i.e.,

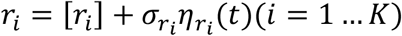

where [*r_i_*] is the mean of *r_i_*, the coefficients *σ_r_i__* is the standard deviation of the perturbations, and the *η_r_i__*(*t*) is a Gaussian white noise with zero mean, time-independent, and statistically independent. Then we could define the strength of extrinsic noise for reaction rates as Fano factor 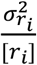. We can extend the chemical Langevin equation in Eq. (2) to include extrinsic fluctuations [7].

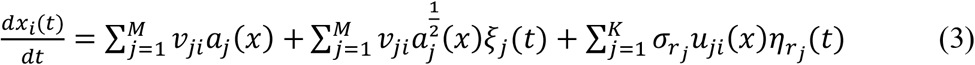

where 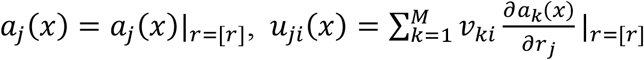.

Eq. (3) described both internal noise and extrinsic noise. The noise of a molecular species *i* is defined as coefficient of variation 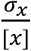. where *x* denote the molecular species and square brackets denote the number of molecular species in the steady state. We could simulate the noise of any molecule species in the system (1) using Gillespie algorithm [8]. In this system, we calculated the noise of the miRNA total abundance (*m + c*) to simulate real sequencing data.

Obviously, when introducing a new target gene (competing RNA) that shares the same regulatory molecule species miRNA, we can extend the ODEs model as follows.

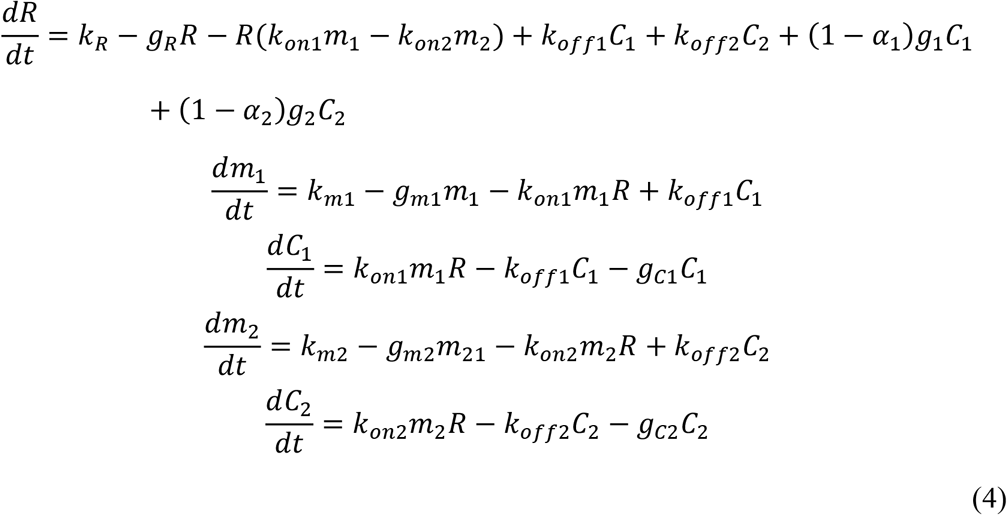

### 3. Influence of the accelerated degradation on microRNA-mediated noise regulation

In order to explore the effect of miRNA accelerated mRNA degradation rate on mRNA noise, we therefore extended the model (1) to allow for variable degrees of accelerated degradation rate Firstly, according to previous studies [5], it is assumed that the association and dissociation of mRNA-microRNA complexes is much faster than the degradation of molecular species. Therefore, based on the quasi-steady state approximation of mRNA-microRNA complexes,

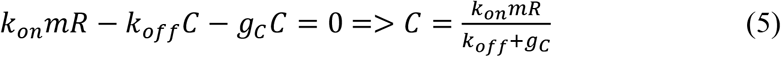

Then the levels of total mRNA *m^T^* can then be expressed as

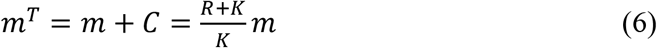

where 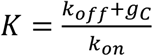 is the Michaelis-Menten constant of the mRNA-microRNA complex. Then the extend model is

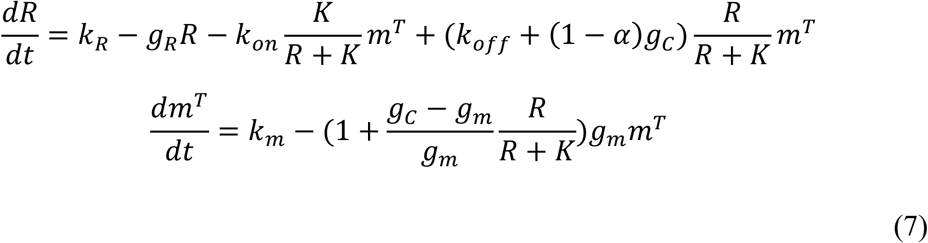

The influence of miRNA accelerated mRNA degradation is thus given by the ratio of microRNA-induced mRNA degradation rate to mRNA degradation rate, ie, 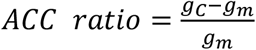. Let the intrinsic Gaussian white noise *ξ_j_*(*t*) described in the supplementary model section (Eq. (2)) as zero, we could simulate the influence of *ACC ratio* on extrinsic noise of mRNA total. The parameters used in simulation are derived from previous researches on gene expression noise regulated by miRNA and competing RNA effects (See Table.S3) [6, 9].

---

**Fig.S1.**
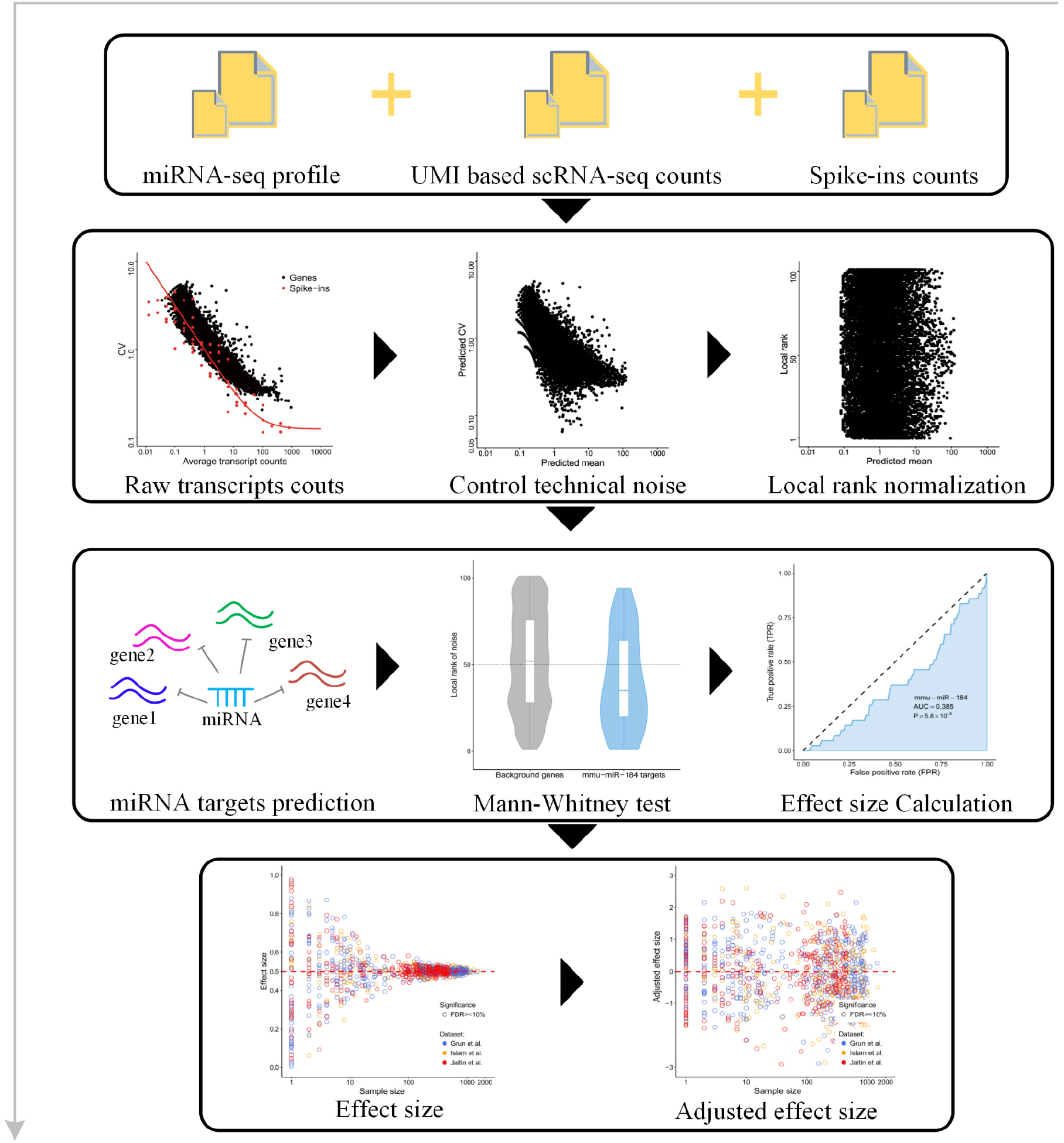
The workflow of gene expression noise analysis.

**Fig.S2.**
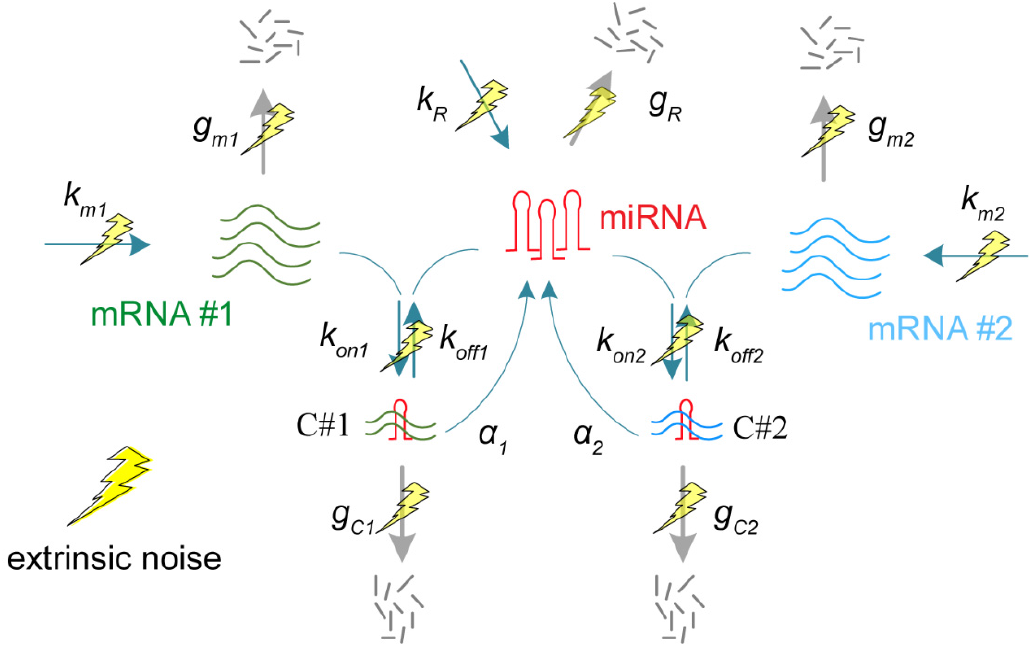
A kinetics model for miRNA regulation. A coarse-gained model for expression noise of a miRNA regulated gene. Noise in mRNA expression originates from stochastic biochemical reactions in the production of the mRNA (intrinsic noise) and fluctuation in the reaction kinetic rates caused by variable external factors in environmental (extrinsic noise; yellow lightning symbols).

**Fig.S3.**
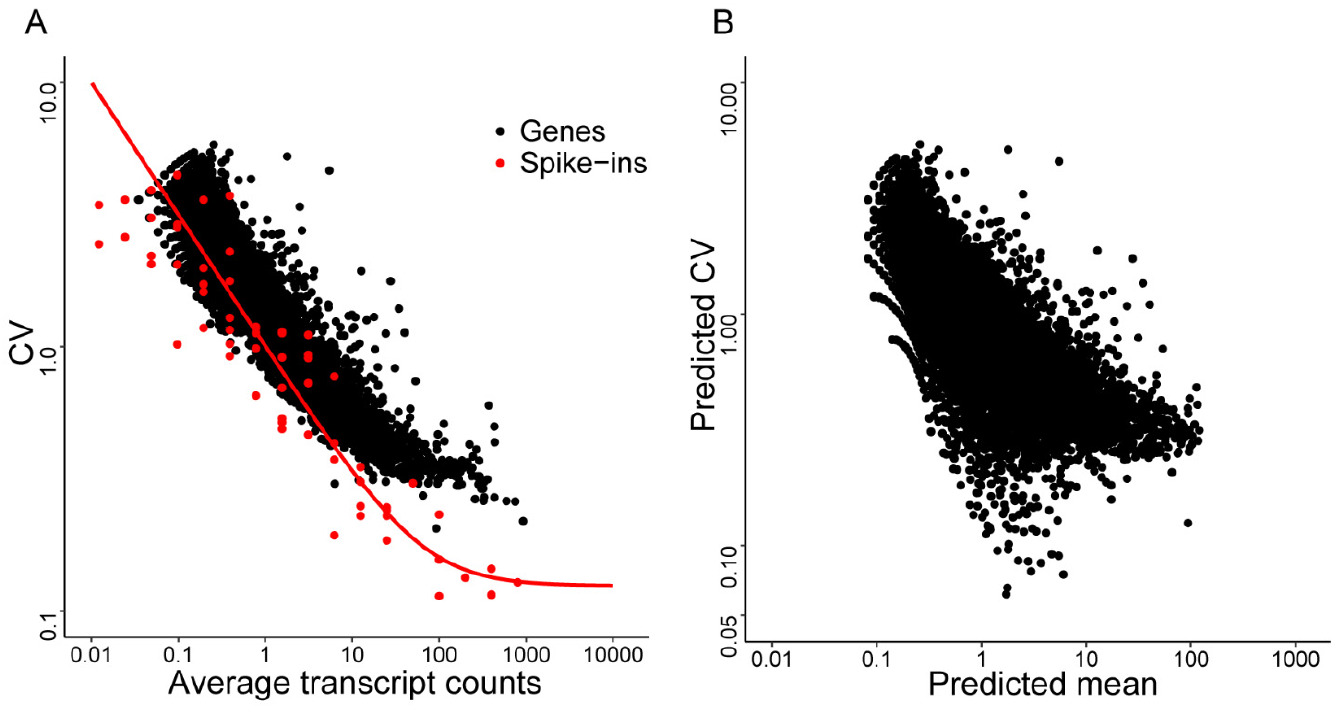
Noise quantification and normalization for mESCs transcripts. (A) Coefficient of variation(CV) versus mean of transcript counts for all detected genes in raw data from scRNA-seq experiment in mouse embryonic stem cells, black dots and red dots indicate true genes and spike-ins respectively. (B) Predicted CV of transcript counts versus predicted mean transcript counts for all detected genes after removing technical noise with the help of spike-ins.

**Fig.S4.**
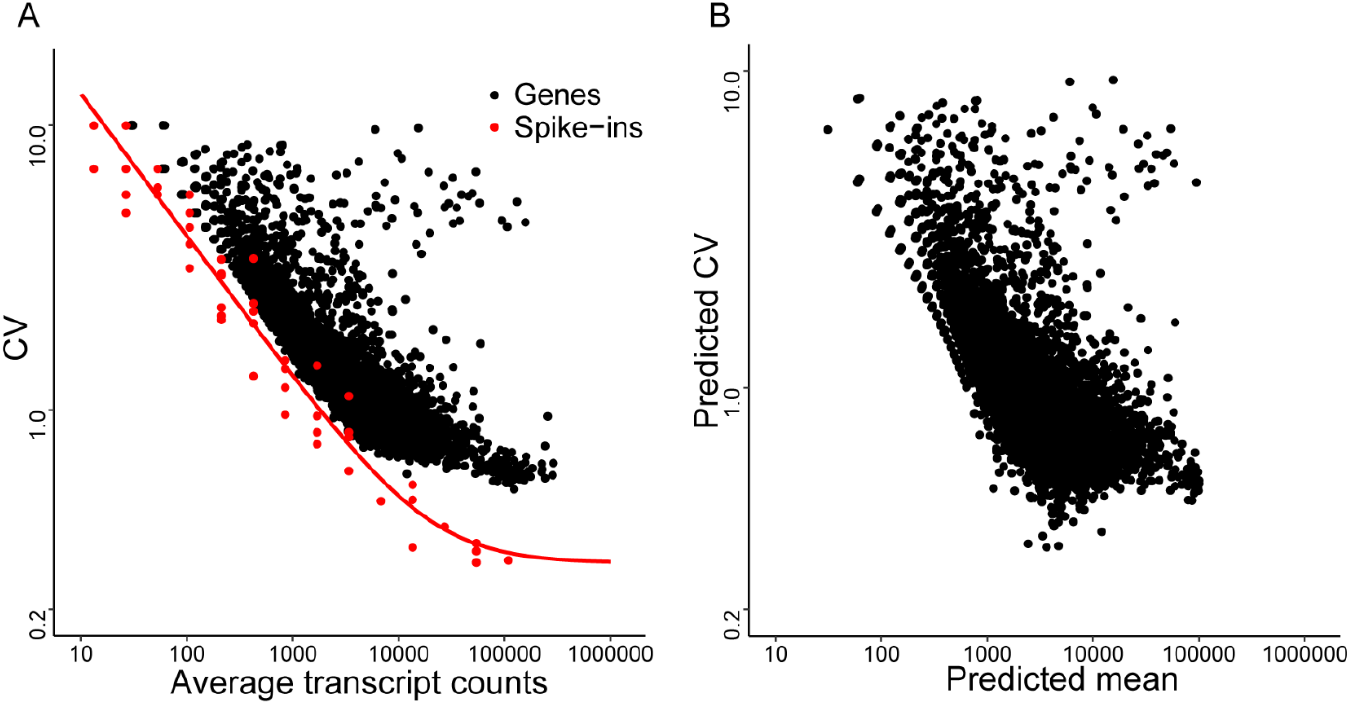
Noise quantification and normalization for mouse intestinal stem cell transcripts. (A) Coefficient of variation(CV) versus mean of transcript counts for all detected genes in raw data from scRNA-seq experiment in mouse intestinal stem cells, black dots and red dots indicate true genes and spike-ins respectively. (B) Predicted CV of transcript counts versus predicted mean transcript counts for all detected genes after removing technical noise with the help of spike-ins.

**Fig.S5.**
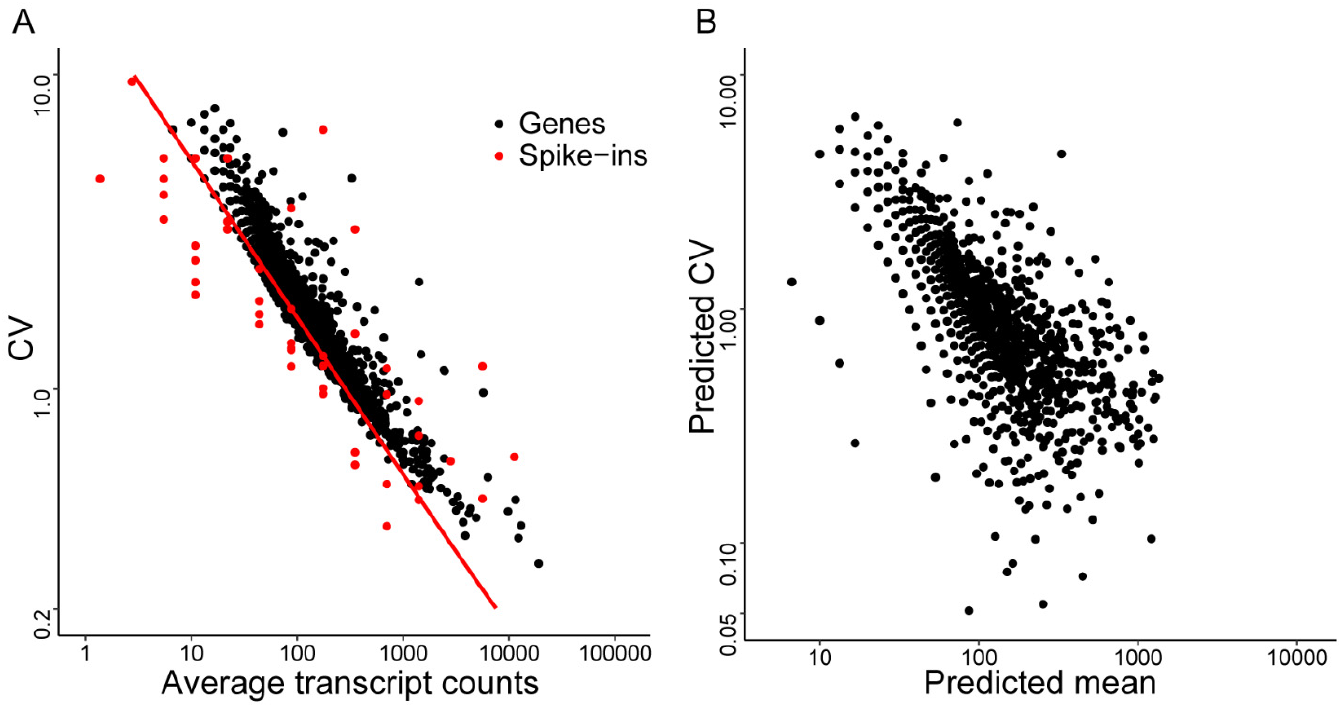
Noise quantification and normalization for mouse dendritic cell transcripts. (A) Coefficient of variation(CV) versus mean of transcript counts for all detected genes in raw data from scRNA-seq experiment in mouse dendritic cells, black dots and red dots indicate true genes and spike-ins respectively. (B) Predicted CV of transcript counts versus predicted mean transcript counts for all detected genes after removing technical noise with the help of spike-ins.

**Fig.S6.**
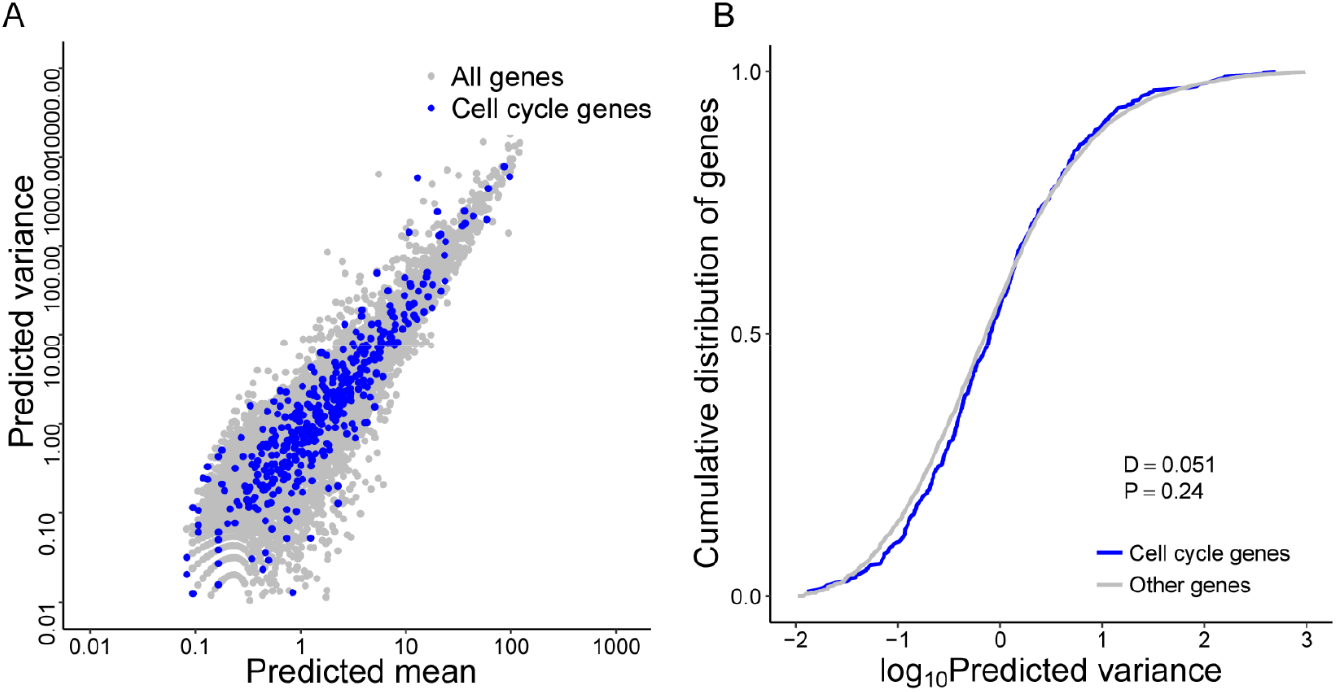
Comparison of variance between cell cycle related genes and other genes for mESCs. (A) Predicted variance of transcript count versus mean transcript counts across the entire detected genes for mESCs. Cell cycle related genes are highlighted as blue circles. This set of genes consist of all genes containing “cell cycle” term in whole Gene Ontology terms of “biological process”. (B) Compared the predicted variance cumulative distribution of cell cycle related genes with other genes, cell cycle related genes do not show increased variability and are thus not to lead to false results in the downstream noise analysis.

**Fig.S7.**
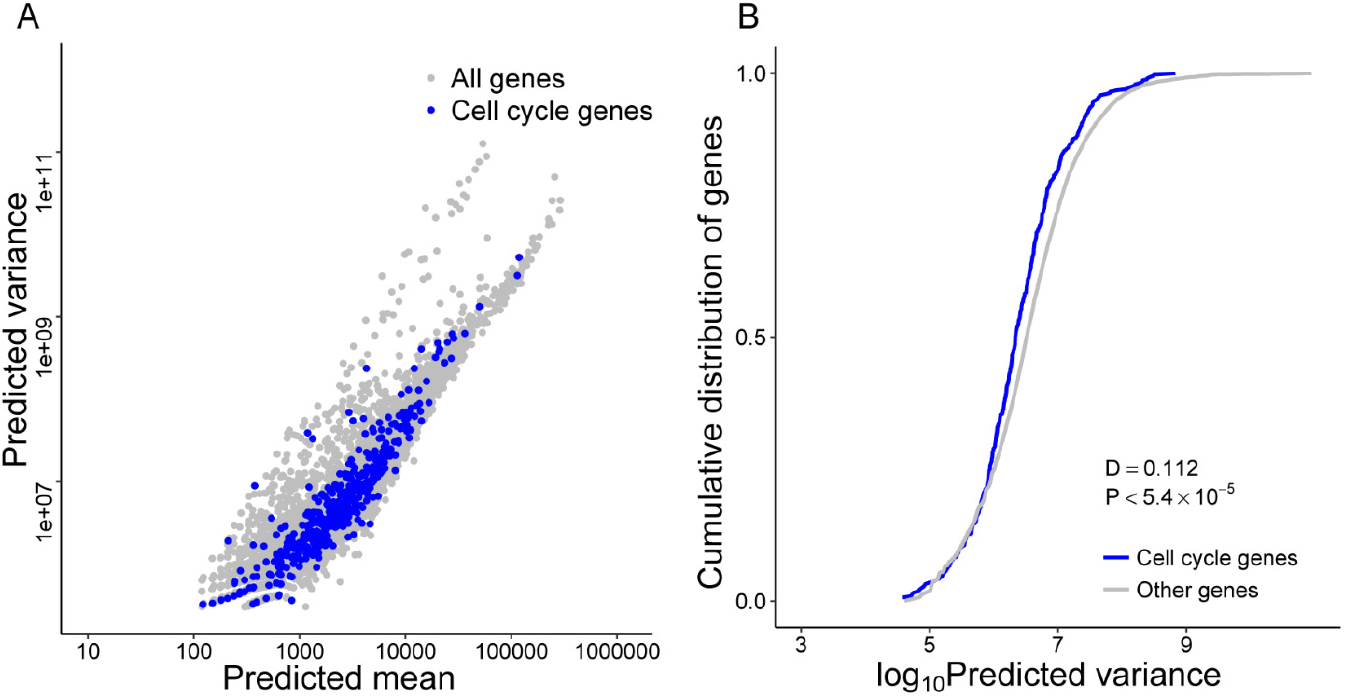
Comparison of variance between cell cycle related genes and other genes for intestinal stem cells. (A) Predicted variance of transcript count versus mean transcript counts across the entire detected genes for intestinal stem cells. Cell cycle related genes are highlighted as blue circles. This set of genes consist of all genes containing “cell cycle” term in whole Gene Ontology terms of “biological process”. (B) Compared the predicted variance cumulative distribution of cell cycle related genes with other genes, cell cycle related genes do not show increased variability and are thus not to lead to false results in the downstream noise analysis.

**Fig.S8.**
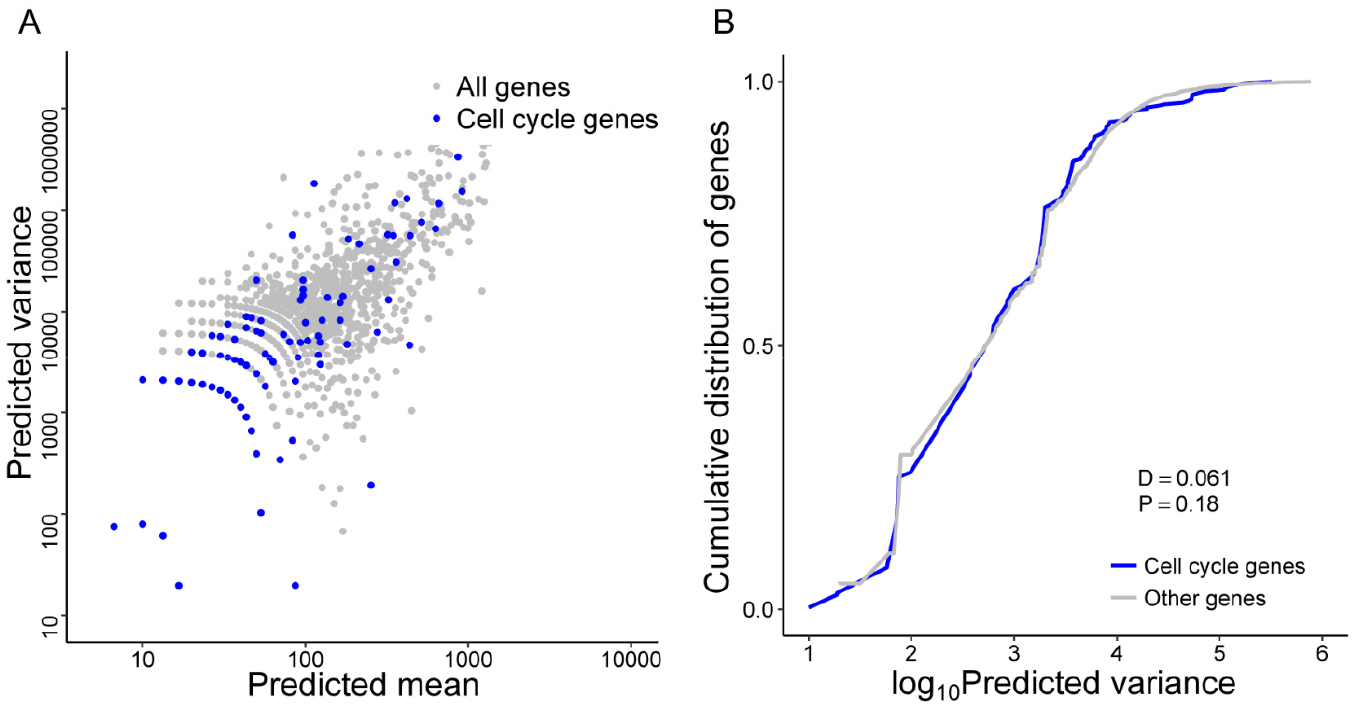
Comparison of variance between cell cycle related genes and other genes for dendritic cells. (A) Predicted variance of transcript count versus mean transcript counts across the entire detected genes for dendritic cells. Cell cycle related genes are highlighted as blue circles. This set of genes consist of all genes containing “cell cycle” term in whole Gene Ontology terms of “biological process”. (B) Compared the predicted variance cumulative distribution of cell cycle related genes with other genes, cell cycle related genes do not show increased variability and are thus not to lead to false results in the downstream noise analysis.

**Fig.S9.**
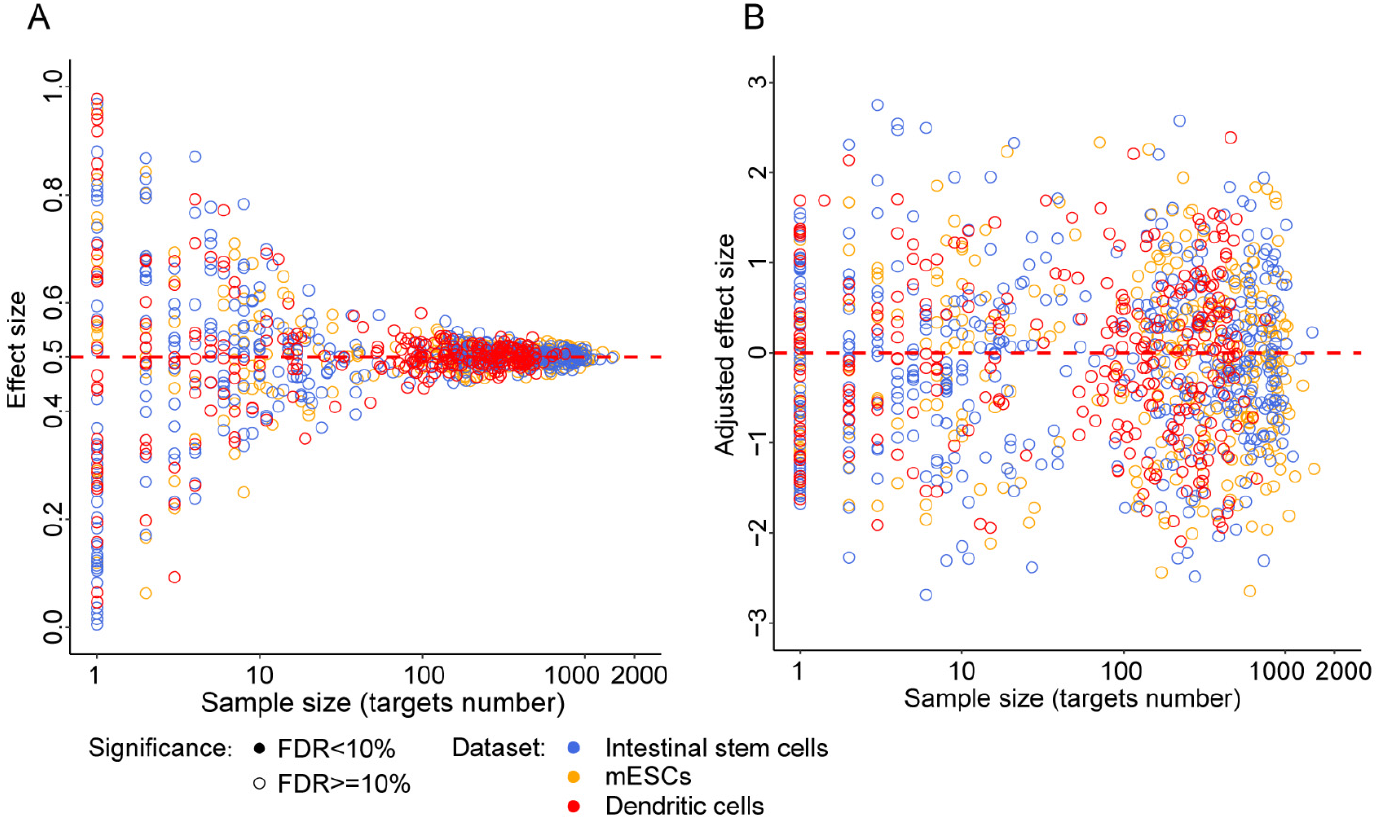
The correlation between effect size and sample size for Mann-Whitney test before correction. (A) Validating the correlation between effect size and test sample size by sampling background samples and variable test samples (x-axis) from all genes randomly in three cell types datasets. The three cell types datasets are highlighted with different color. And all tests are not significant in random experiment (FDR<10%). This simulation result show that effect size is a strongly correlated with the test sample size. (B). An additional effect size correction step to remove the effects of sample size by calculating the adjust effect size as z-score according the distribution of effect size. This plot shows the adjust effect size after correction.

**Fig.S10.**
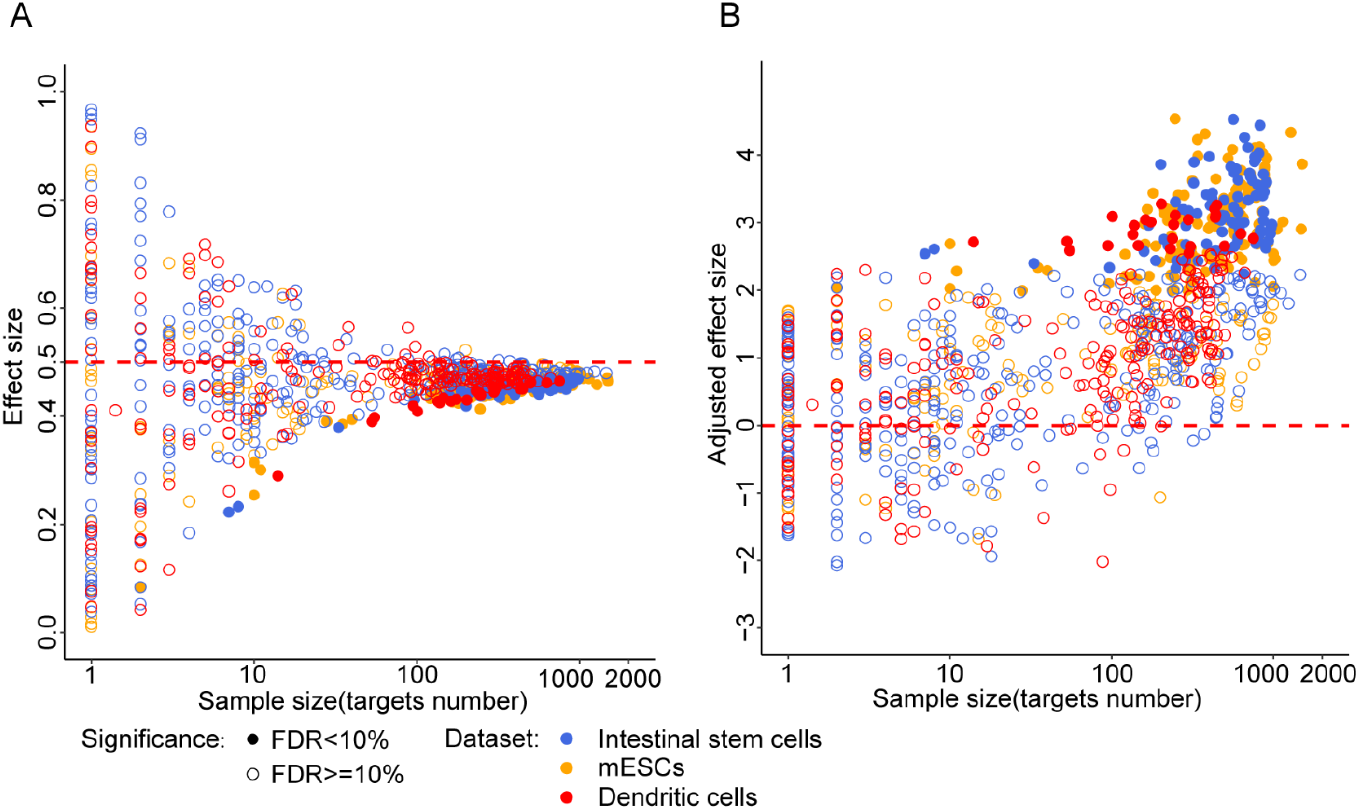
The correlation between effect size and sample size for Mann-Whitney test after correction. (A) Effect size vesus number of target (test sample size) for real data. This results shows that there is a noise biases for all three cell line datasets. Effect size less than 0.5 indicates that local rank CV of miRNA target genes are smaller than non-targets. (B). An additional effect size correction step for real data.

**Fig.S11.**
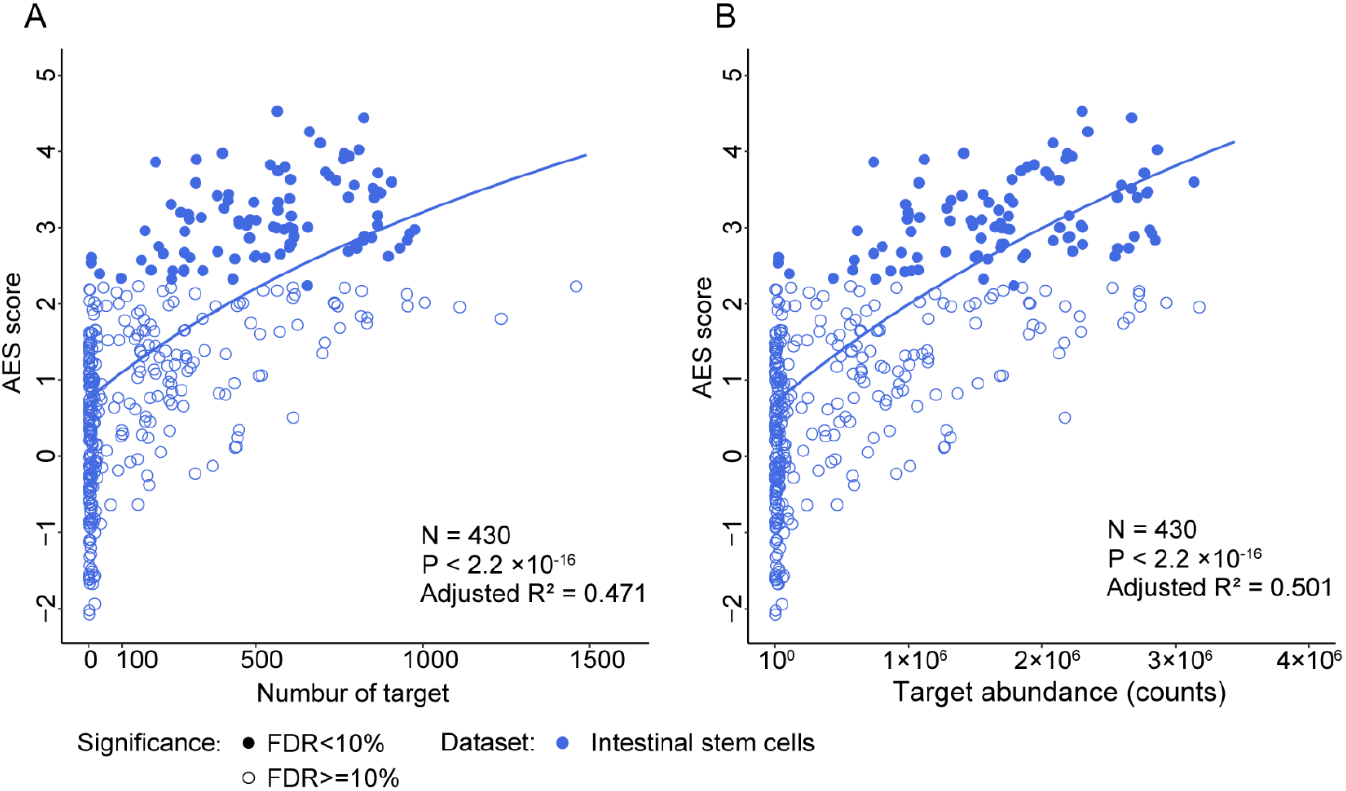
miRNAs with large target pool associated with better noise suppression for intestinal stem cells. (A) AES score of Mann-Whitney test versus number of targets across all detected miRNA in intestinal stem cells. Curves were fit to *a + b*log_10_(*x + c*) where a, b and c were determined by least squares error. Fitted parameters are shown in Table.S1. (B) Similar fitting curve to those shown in A for targets pool abundance.

**Fig.S12.**
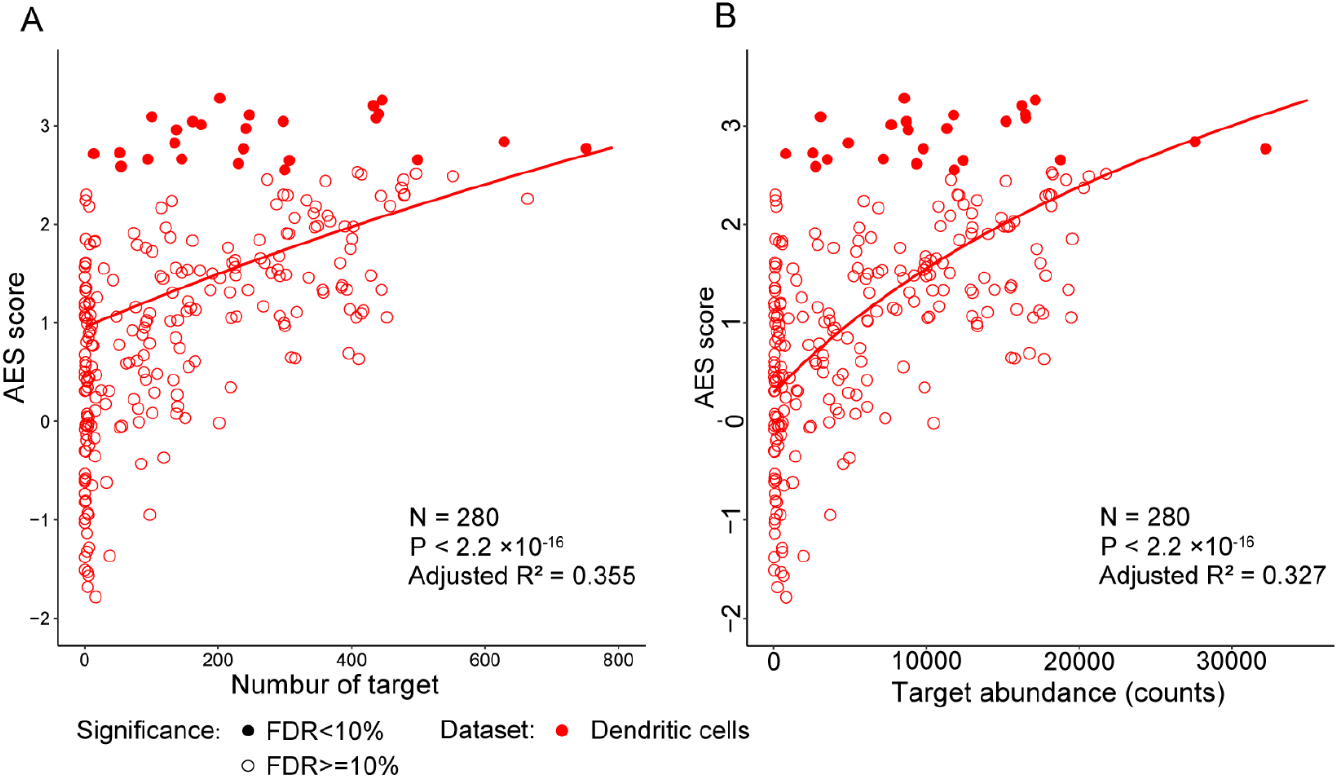
miRNAs with large target pool associated with better noise suppression for dendritic cells. (A) AES score of Mann-Whitney test versus number of targets across all detected miRNA in dendritic cells. Curves were fit to *a + b*log_10_(*x + c*) where a, b and c were determined by least squares error. Fitted parameters are shown in Table.S1. (B) Similar fitting curve to those shown in A for targets pool abundance.

**Fig.S13.**
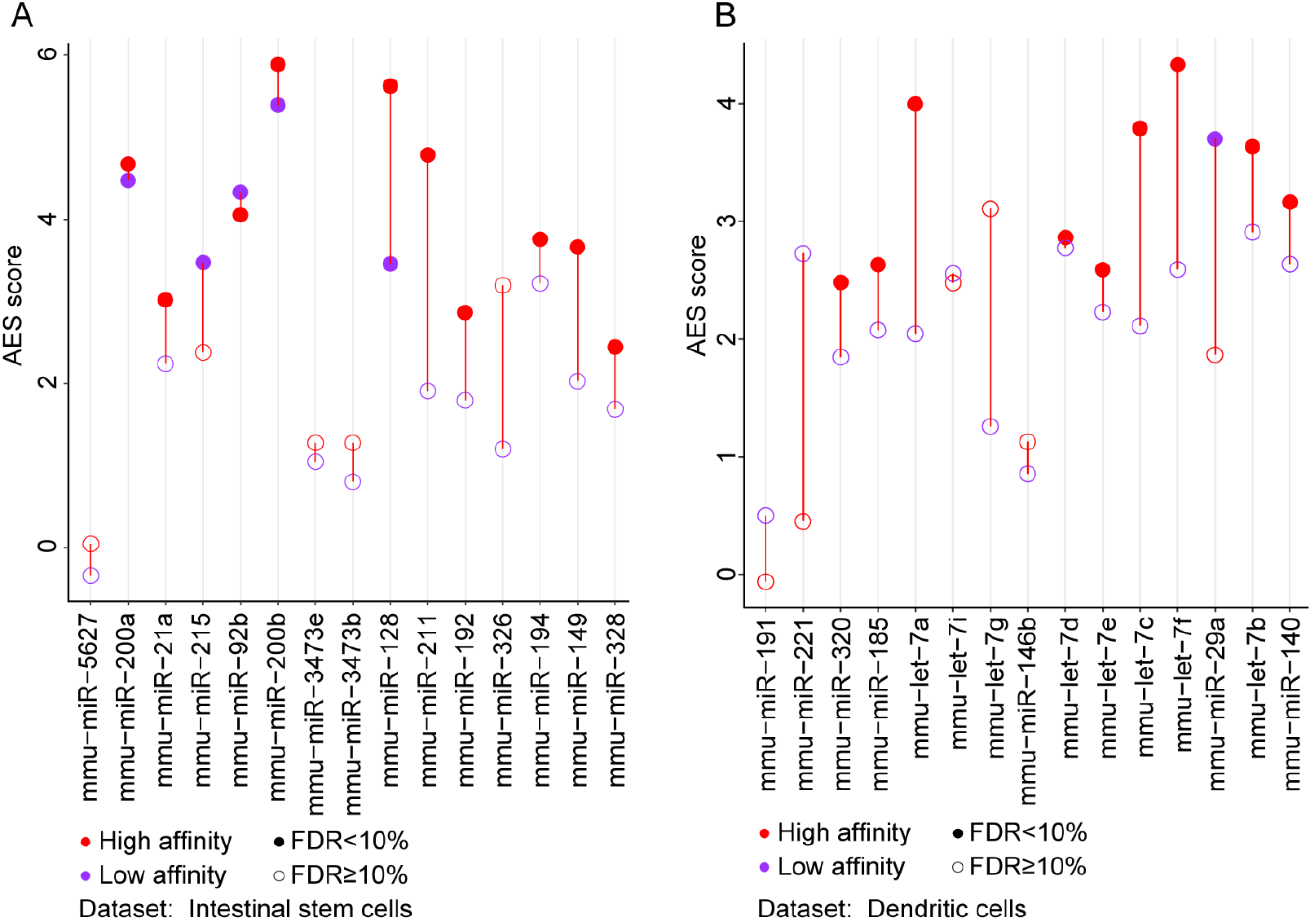
Strong interactions will enhance the effect of noise reduction. (A-B) Noise disparity between the target genes of Top15 abundant miRNA for strong (red) and weak interaction strength (purple) in intestinal stem cells (left) and dendritic cells (right) respectively.

**Fig.S14.**
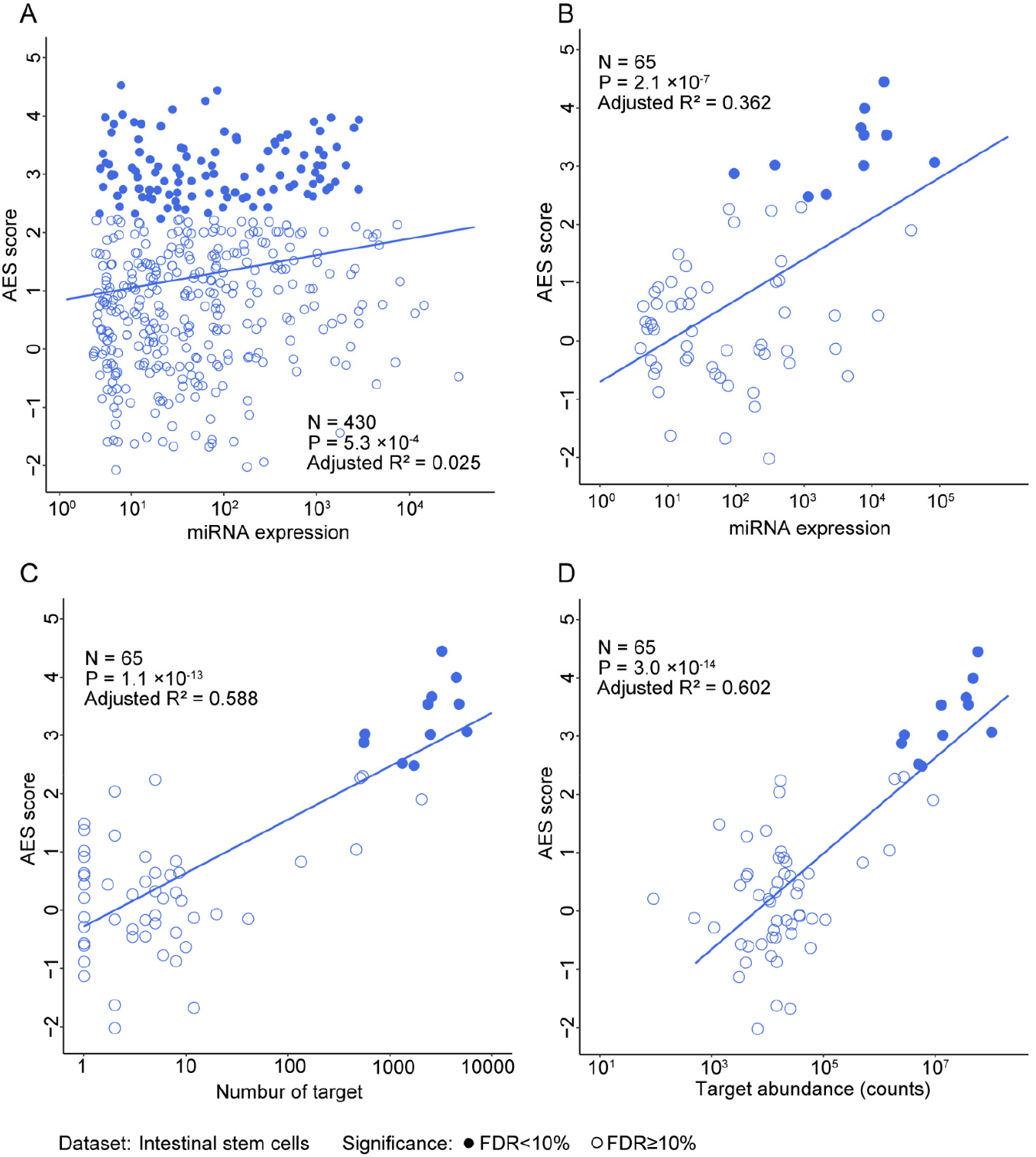
Gene expression noise are repressed by miRNA co-regulation sub-network for intestinal stem cells. (A-B) AES score of Mann-Whitney test versus miRNA expression for miRNA individuals and subnetwork respectively in intestinal stem cells. Curves were fit to *a + b*log_10_(*x + c*) where a and b were determined by least squares error. Fitted parameters are shown in Table.S2. (C) AES score of Mann-Whitney test after sub-network clustering versus number of targets across all cluster miRNA in intestinal stem cells. (D) Similar fitting curve to those shown in C for targets pool abundance.

**Fig.S15.**
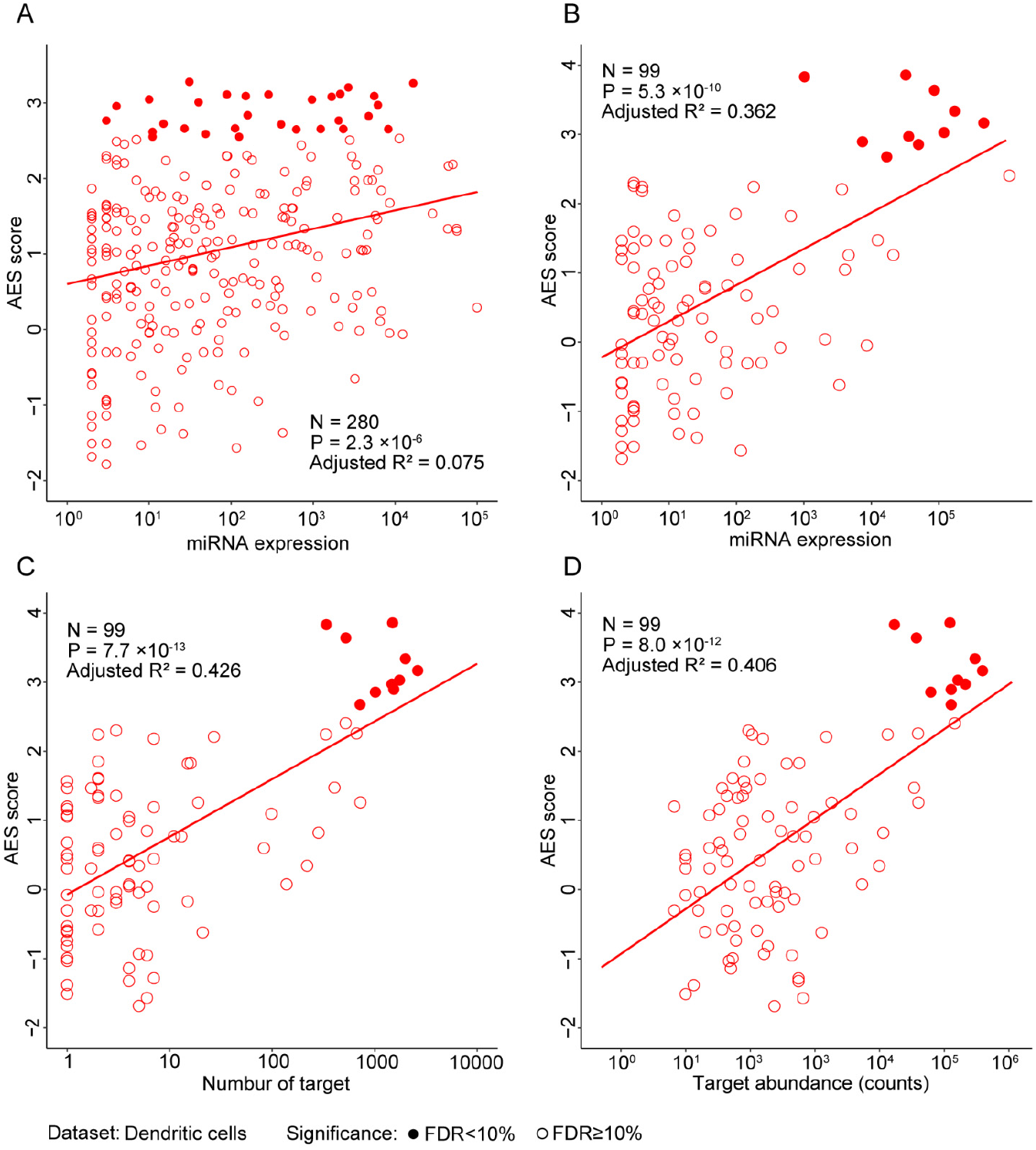
Gene expression noise are repressed by miRNA co-regulation sub-network for dendritic cells. (A-B) AES score of Mann-Whitney test versus miRNA expression for miRNA individuals and sub-network respectively in dendritic cells. Curves were fit to *a + b*log_10_ *x* where a and b were determined by least squares error. Fitted parameters are shown in Table.S2. (C) AES score of Mann-Whitney test after sub-network clustering versus number of targets across all cluster miRNA in dendritic cells. (D) Similar fitting curve to those shown in C for targets pool abundance.

**Fig.S16.**
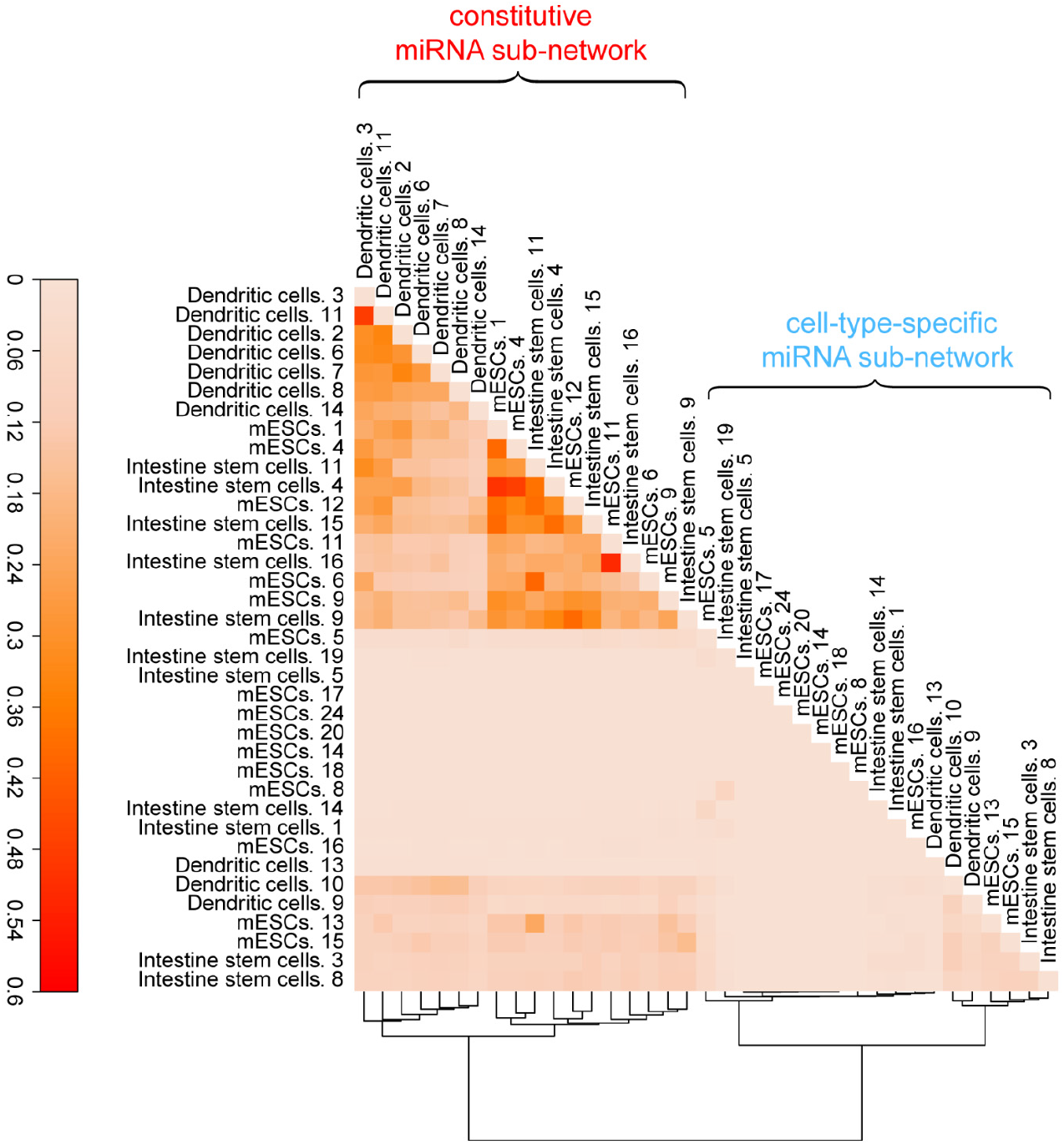
Similarity matrix for all miRNA sub-networks from three cell types, Sub-networks were divided into constitutive sub-networks and cell-type-specific sub-networks by hierarchical clustering of similarity matrix. Sub-network similarity matrix is calculated using Jaccard similarity of their target genes. The numbers in network name indicates the sub-networks ID. Details miRNAs of sub-network are shown in Table. S4.

**Fig.S17.**
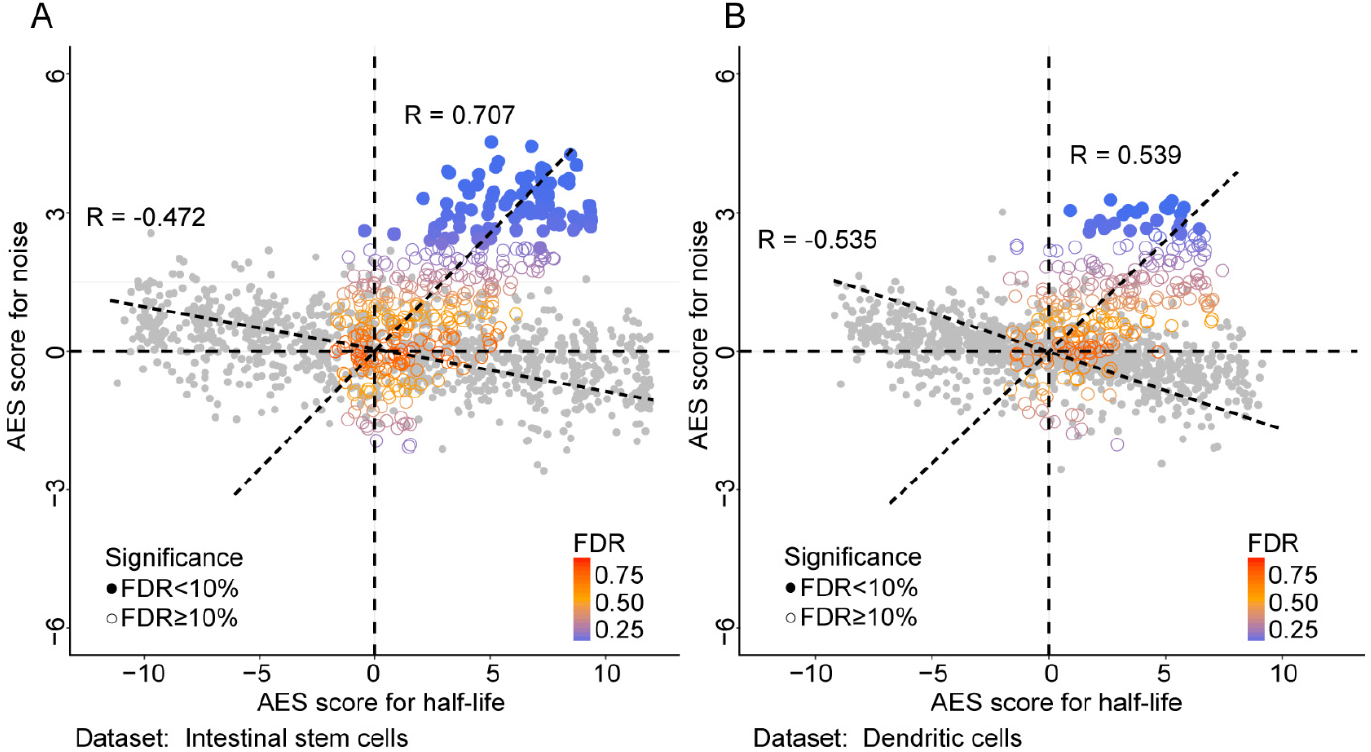
The correlation between noise and mRNA half-life. (A-B) Scatterplot of gene expression noise AES score versus mRNA half-life AES score for random gene sets of varying sample size (200–2000) by weighted sampling to half-life of mRNA (grey) and real miRNA target gene set (blue) for intestinal stem cells and dendritic cells datasets respectively. (Fig.15A, grey, Spearman correlation = −0.472, color, Spearman correlation = 0.707; Fig.15B, grey, Spearman correlation = −0.535, color, Spearman correlation = 0.539)

**Fig.S18.**
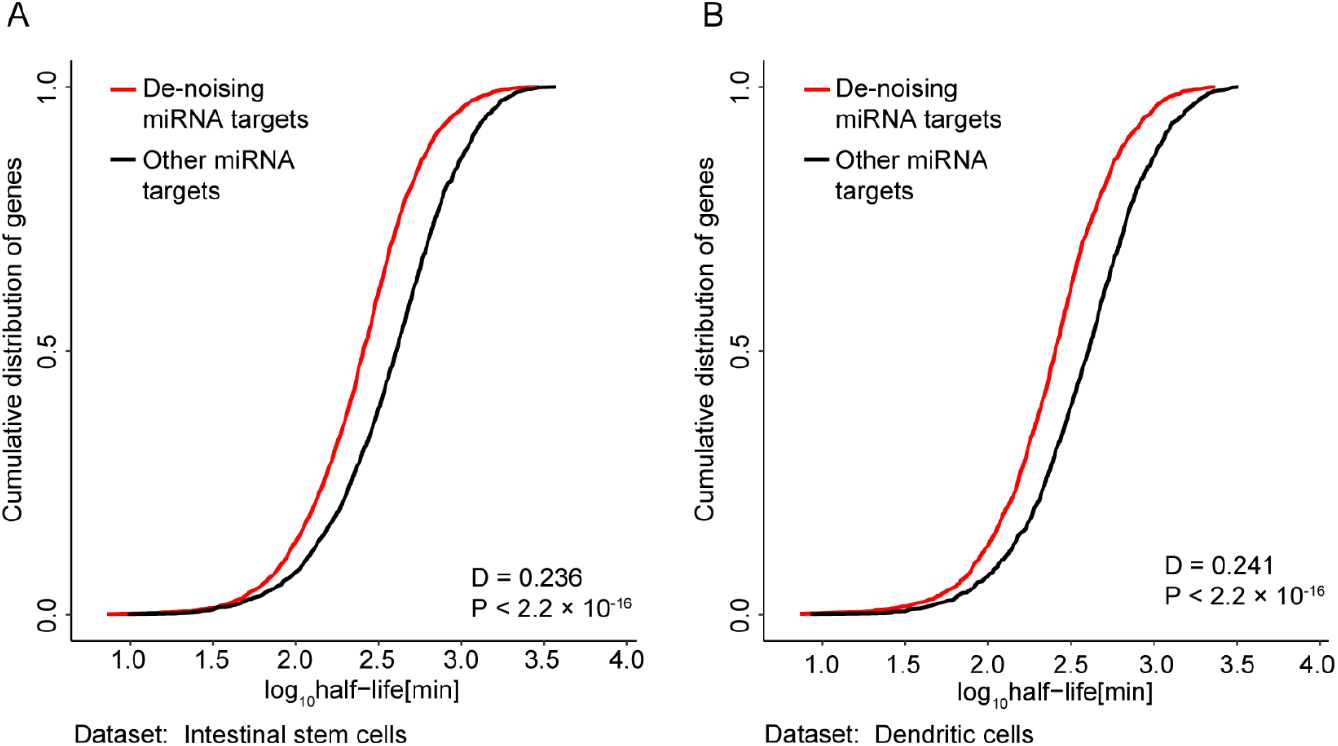
The distribution of mRNA half-life time for miRNA targets. (A-B) Compared the mean mRNA half-lives cumulative distribution of de-noising miRNAs target genes with target genes of remain miRNAs in intestinal stem cells and dendritic cells respectively. We observe that the targets of de-noising miRNAs have much shorter half-life than those targets of remain miRNAs.

Information of parameters used in this study

**Table S1.** Fitted parameters in this article

|  | a | b | c |
| --- | --- | --- | --- |
| Fig.2A | -12.300 | 4.899 | 633.0 |
| Fig.2B | -10.794 | 3.258 | 4733.0 |
| Fig.4B | 1.005 | 0.440 | 0.0 |
| Fig.3C | -19.729 | 6.828 | 1132.828 |
| Fig.3D | -8.549 | 4.266 | 73.157 |
| Fig.S11A | -21.641 | 7.573 | 907.0 |
| Fig.S11B | -66.125 | 10.318 | 3000000 |
| Fig.S14A | 1.658 | 0.530 | 0.0 |
| Fig.S12A | -29.697 | 9.689 | 1457.0 |
| Fig.S12B | -23.425 | 5.679 | 15000.0 |
| Fig.S15A | 0.602 | 0.244 | 0.0 |

**Table S2.** Fitted parameters in this article

|  | a | b |
| --- | --- | --- |
| Fig.4D | -0.001 | 1.137 |
| Fig.4E | -0.884 | 0.908 |
| Fig.4C | -0.803 | 0.939 |
| Fig.S14C | -0.278 | 0.915 |
| Fig.S14D | -3.115 | 0.820 |
| Fig.S14B | -0.703 | 0.702 |
| Fig.S15C | -0.072 | 0.835 |
| Fig.S15D | -0.929 | 0.650 |
| Fig.S15E | -0.222 | 0.523 |

**Table S3.** Parameters in physical model

| Parameter | value | Parameter | value |
| --- | --- | --- | --- |
| $k_R$ | 3 | $k_{m2}$ | 2 |
| $g_R$ | 1/30 | $g_{m2}$ | 1/2 |
| $k_{m1}$ | 2 | $k_{on2}$ | 2 |
| $g_{m1}$ | 1/2 | $k_{off2}$ | 2 |
| $k_{on1}$ | 2 | $g_{c1}$ | 1 |
| $k_{off1}$ | 1 | $g_{c2}$ | 1 |
| $\alpha_1$ | 0.5 | $\alpha_2$ | 0.5 |
| Fano factor for all parameters |  | 0.25 |  |

Significantly de-noising miRNA sub-networks

**Table S4.**
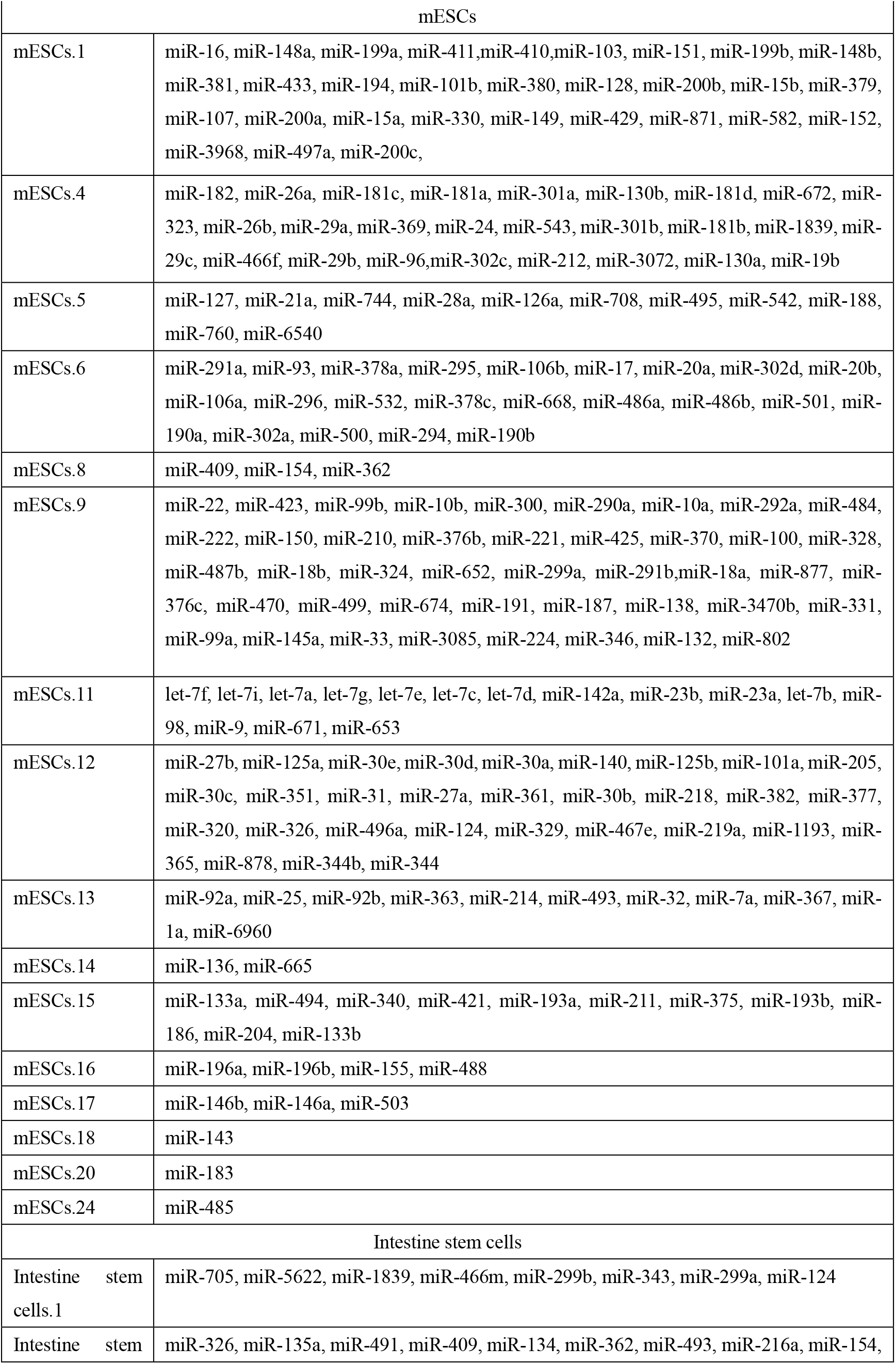
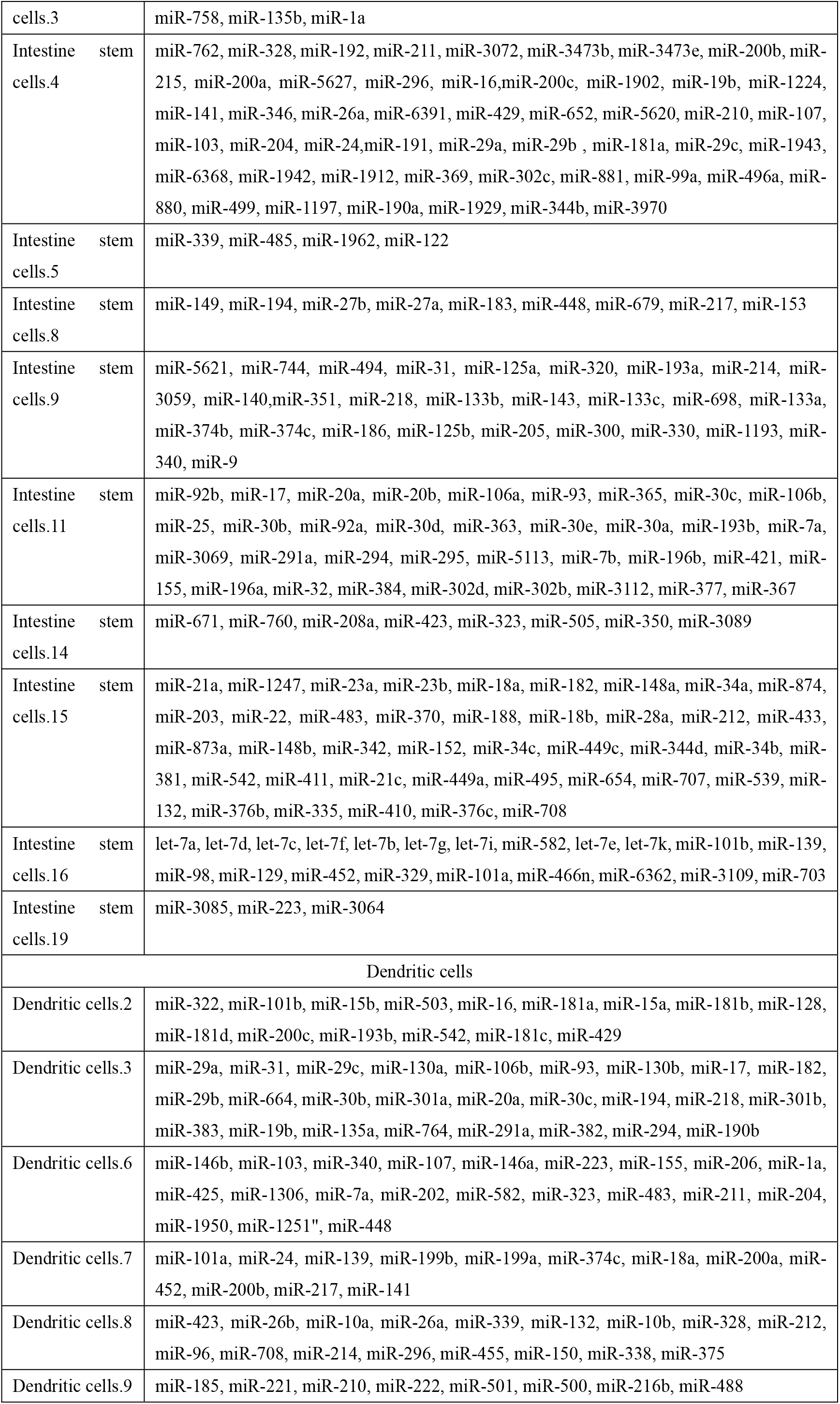
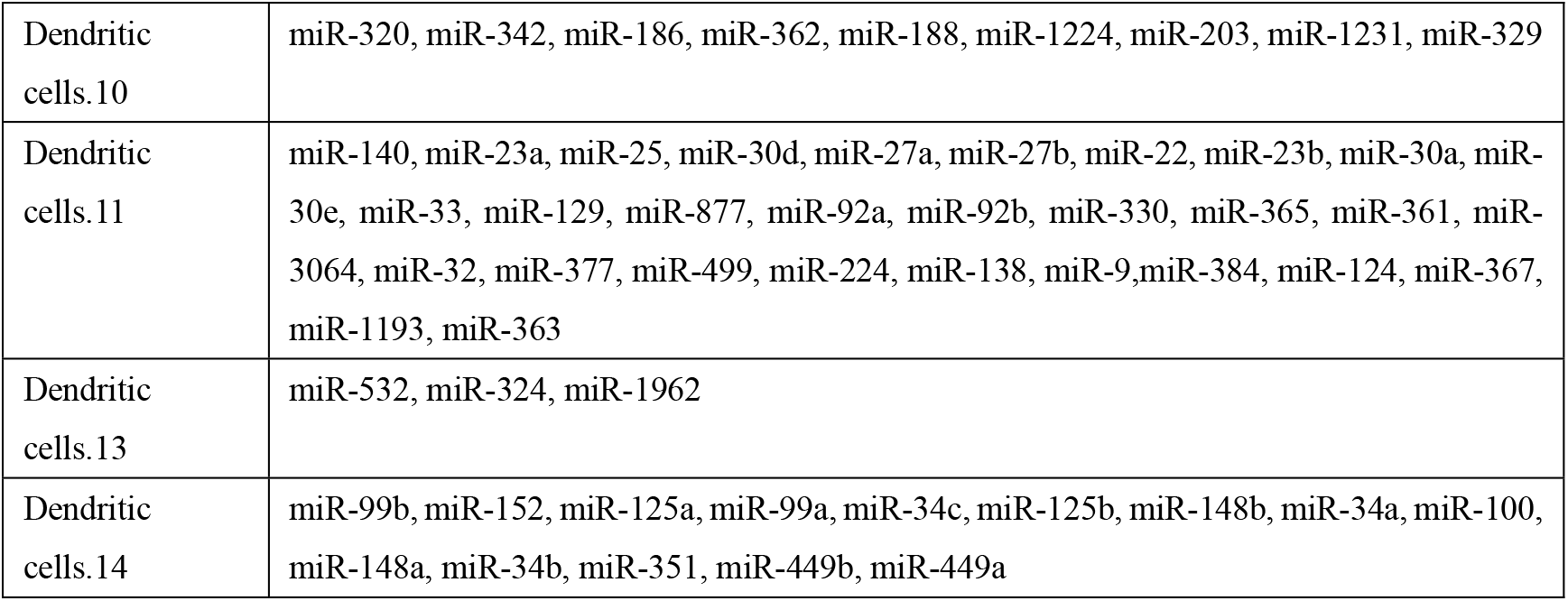
Significantly de-noising miRNA sub-networks for three cell types

